# The obligate intracellular bacterium *Orientia tsutsugamushi* differentiates into a developmentally distinct extracellular state

**DOI:** 10.1101/2020.03.03.975011

**Authors:** Sharanjeet Atwal, Jantana Wongsantichon, Suparat Giengkam, Kittirat Saharat, Yanin Jaiyen Pittayasathornthun, Suthida Chuenklin, Loo Chien Wang, Taerin Chung, Hyun Huh, Sang-Hyuk Lee, Radoslaw M. Sobota, Jeanne Salje

## Abstract

*Orientia tsutsugamushi* (Ot) is an obligate intracellular bacterium in the family Rickettsiaceae that causes scrub typhus, a severe mite-borne human disease. Its mechanism of cell exit is unusual amongst Rickettsiaceae, as Ot buds off the surface of infected cells enveloped in plasma membrane. Here, we show that Ot bacteria that have budded out of host cells are in a distinct developmental stage compared with intracellular bacteria. We refer to these two stages as intracellular and extracellular bacteria (IB and EB, respectively). These two forms differ in physical properties: IB is elongated, and EB is round. Additionally, IB has higher levels of peptidoglycan and is physically robust compared with EB. The two bacterial forms differentially express proteins involved in bacterial physiology and host-pathogen interactions, specifically those involved in bacterial dormancy and stress response, secreted bacterial effectors, and outer membrane autotransporter proteins ScaA and ScaC. Whilst both populations are infectious, entry of IB Ot is sensitive to inhibitors of both clathrin-mediated endocytosis and macropinocytosis, whereas entry of EB Ot is only sensitive to a macropinocytosis inhibitor. Our identification and detailed characterization of two developmental forms of Ot significantly advances our understanding of the intracellular lifecycle of an important human pathogen.

**Author Summary:** *Orientia tsutsugamushi* (Ot) is a bacterial pathogen that causes scrub typhus, a mite-transmitted human disease. This illness is traditionally known to be endemic in the Asia-Pacific, but recent reports of *Orientia-*like organisms from the Middle East, Africa, and Latin America suggest that it may be globally distributed. Scrub typhus is associated with high mortality if not treated promptly with appropriate antibiotics. Ot is a highly specialized bacterium that can only replicate within living cells, either within the mite vector or in mammalian or human hosts. Ot exits infected cells using a unique mechanism that involves budding off the surface of infected cells. We have discovered that this unusual aspect of its lifecycle involves the bacteria themselves differentiating into a distinct growth form. Different growth forms have not been described in other members of the family Rickettsiaceae, and no other family members have been shown to bud out of host cells in a manner similar to Ot. We find that the two forms of Ot, which we refer to as intracellular and extracellular bacteria (IB and EB respectively), differ in physical properties and protein expression and infect cells through different mechanisms. The identification of structurally and functionally distinct forms of Ot elucidates a vital aspect of this pathogen’s intracellular life cycle. The two forms are likely to have different antibiotic susceptibilities, therefore our findings may advance the development of novel interventions aimed at inhibiting Ot growth in scrub typhus patients.

## Main

*Orientia tsutsugamushi* (Ot) is an obligate intracellular Gram-negative bacterium that causes the mite-transmitted human disease scrub typhus. This condition is one of the most severe rickettsial diseases; it is associated with a median 6% mortality in untreated patients^1^. Scrub typhus is a leading cause of severe febrile illness in the Asia-Pacific, a region containing two-thirds of the world’s population. Recent reports from Latin America, the Middle East, and Africa suggest that the disease is more globally distributed^2^. Knowledge of the fundamental biology of this organism lags behind that of pathogens of equivalent prevalence and severity. Historical underestimations of Ot incidence, its classification as a biosafety level 3 pathogen, its fastidious growth in cultured cells, and the fact that it remains genetically intractable^3^ likely account for the paucity of studies focused on this pathogen.

The order Rickettsiales contains two major families: Anaplasmataceae and Rickettsiaceae^4^. Members of the Anaplasmataceae, which include *Anaplasma* spp. and *Ehrlichia* spp., reside within membrane-bound vacuoles inside host cells. The developmental cycle of certain Anaplasmataceae family members is biphasic and involves switching between a non-infectious intracellular replicative form (reticulate body) and an infectious extracellular form (elementary body). While members of the Rickettsiaceae, namely *Rickettsia* spp. and Ot, do not appear to differentiate into distinct forms during their intracellular lifecycle, Ot is unique amongst the characterized Rickettsiaceae in budding off the surface of infected host cells in a manner reminiscent of enveloped viruses^5^. This observation led us to investigate whether budded Ot is in a distinct developmental stage compared with intracellular counterparts.

Ot has been shown to enter host cells using the zipper-like invasion mechanism, which is common to many Rickettsiales members^4^. This process is initiated by binding of bacterial surface adhesins to receptors expressed on mammalian cells. Adhesins such as TSA56, ScaA, and ScaC^6-8^ bind to host cell fibronectin and syndecan-4; these proteins then interact with integrin α5ß1, leading to activation of intracellular signaling that culminates in actin remodeling and clathrin-mediated endocytosis^9-11^. It is important to emphasize that these studies were entirely focused on intracellular Ot, as they were performed on bacteria isolated from host cells by mechanical lysis of infected cells. However, the re-entry mechanism utilized by bacteria that have budded out of cells may be influenced by the presence of an enveloping host cell membrane and/or its differentiation into a distinct state.

Here, we show that Ot exists in distinct intracellular (IB) and extracellular (EB) bacterial populations. We report the characterization of these two states in terms of their physical properties, metabolic activity, and gene expression profiles and demonstrate that they enter host cells using different mechanisms.

## Results

### IB and EB forms of Ot differ in physical properties and metabolic activity

Ot is typically described as a coccobacillary bacterium. Its small size and propensity to aggregate make it difficult to obtain clear images of this bacterium using conventional light microscopy. To circumvent this challenge, we labelled Ot with an antibody against the abundant surface protein TSA56 and employed stochastic optical reconstruction microscopy (STORM) for high-resolution imaging. We first investigated the shape of bacteria located inside and outside host cells (Fig. 1A, B and Supp. Fig. 1). Since Ot begins to appear in the supernatant of infected L929 cells at high levels five days post infection (Supp. Fig. 2) we focused on the shape of EB Ot beginning five days post infection, whilst we measured the shape of IB Ot throughout the infection cycle. We found that EBs were approximately round in shape with a diameter of about 1.2 µm, and they were often aggregated into clusters. In contrast, IBs adopted diverse round and elongated shapes and varied in dimensions; some bacteria were 3-4 µm in length. Additionally, IB size varied during the course of infection. Initially, and up to one-day post-infection, IBs were similar in length to EBs. At later stages, *i.e.,* 2-7 days following infection, IBs were characterized by longer lengths. These kinetic changes in shape correlated with Ot growth dynamics whereby replication starts 24-48 hours post infection^12^. These differences in shape provide evidence for the existence of two separate bacterial forms.

**Fig. 1:**
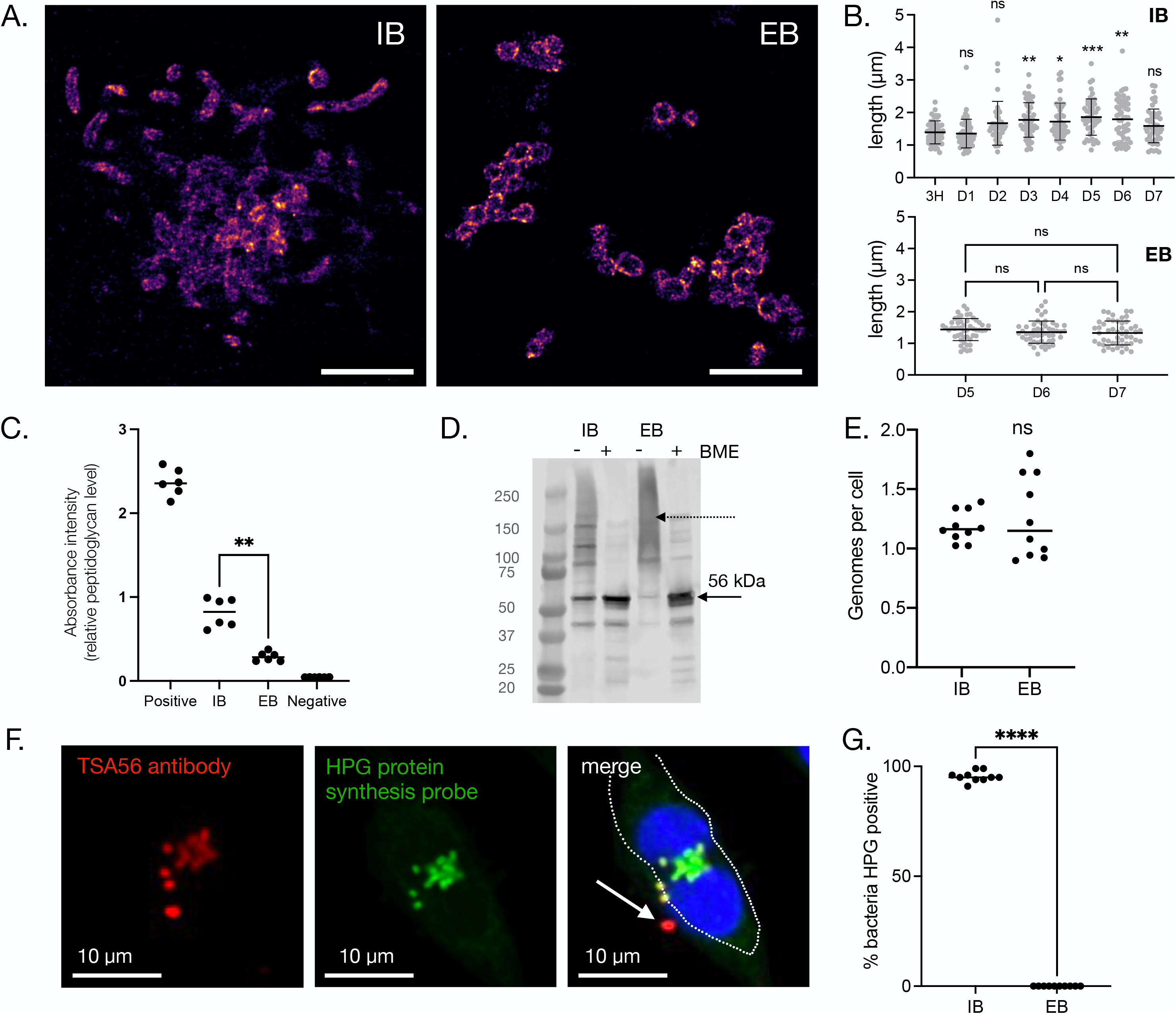
Intracellular and extracellular forms of Ot are physically and metabolically distinct. A. Representative STORM images of IB (5 d.p.i.) and EB (7 d.p.i.) Ot in L929 cells. Bacteria were labelled with a monoclonal antibody against surface protein TSA56 (magenta). Additional images including DIC images of host cells are provided in Supp. Fig. 1. Scale bar = 5 µm. B. Length quantification of individual IB and EB Ot as imaged by STORM. H = hours, D = days post infection. The lengths of 50 individual IB (defined as colocalising with host cells) and EB form (defined as not colocalising with host cells) Ot were measured. Statistical significance was determined using a one-way ANOVA followed by Dunnett’s multiple comparisons. C. Quantification of relative peptidoglycan levels through activation of NOD1 reporter cells, as measured by secreted alkaline phosphatase activity. EB Ot induced less activation than IB, indicating lower levels of peptidoglycan. Identical amounts of IB and EB Ot were used, taken from L929 cells 5 days post infection. The results of six independent replicates, carried out on two different days, are shown and the statistical significance was determined using a Mann-Whitney test. Positive control = meso-diaminopimelic acid, negative control = PBS. D. Immunoblot showing the formation of disulfide cross-linked aggregates of the surface protein TSA56, observed in the absence of the reducing agent beta-mercaptoethanol (BME). This effect was stronger in EB compared with IB Ot, taken five days post infection from L929 cells. The expected molecular weight of TSA56 (56 kDa) is indicated with an arrow, and a dashed arrow depicts higher-order aggregates. E. Genome copy number per cell as measured by qPCR; bacterial cell number was determined by immunofluorescence microscopy. Quantification of 10 individual imaging fields is shown, and statistical significance determined using a Mann-Whitney test. F. Confocal microscopy image of an L929 cell infected with Ot for five days and labelled with the clickable methionine analog homopropargylglycine (HPG, green) and an antibody against surface protein TSA56 (red). DNA (Hoechst) is shown in blue. The host cell outline is showed by a dashed line, and an arrow indicates a metabolically inactive bacterium located outside the host cell. G. Quantification of metabolic activity, as measured by a detectable HPG signal, in IB and EB Ot. L929 cells were infected with Ot for five days, fixed, and labelled with TSA56 and HPG. Bacteria were identified as being IB or EB, based their position relative to host cells, and the percentage of each population that was HPG-positive was quantified. The percentage positivity for ten randomly imaged field is shown here, and this data is representative of >3 independent repeats of this experiment. Groups were compared using a Mann-Whitney test. Statistical significance for all data in this figure was determined using Graphpad Prism software. * p-value ≤ 0.05, ** p-value ≤ 0.01, *** p-value ≤ 0.001, **** p-value ≤ 0.0001, ns = not significant.

We next investigated potential differences in bacterial cell wall composition. We previously showed that Ot builds a low-abundance peptidoglycan-like structure^13^. Thus, we assessed relative peptidoglycan levels in IB and EB Ot forms using a NOD-1 reporter assay (Fig. 1C). We detected significantly more peptidoglycan in IB cells compared with EB cells. This finding may account for the observed differences in shape (Fig. 1A) as peptidoglycan is required to build elongated cells.

In related structural studies, we capitalized on our previous finding that the abundant surface protein TSA56 is involved in forming disulfide bonds with itself or other proteins, leading to the formation of aggregates^13^. These complexes can be identified as smears following sample preparation in the absence of reducing agents and evaluation by immunoblot analyses. In this study, we found evidence for substantially higher disulfide bond formation in EB Ot than IB Ot, as determined by the presence of a high molecular weight smear (Fig. 1D). We hypothesize that this crosslinked structure confers EB Ot with additional structural rigidity in the absence of peptidoglycan. These experiments were performed in the presence of iodoacetamide, an agent that covalently modifies cysteine residues and prevents them from forming new bonds, thus ensuring that the cross-links were not formed during sample preparation and processing.

Our next goal was to assess potential changes in genome copy number as, during rapid growth, bacteria such as *E. coli* carry multiple genome copies within a single cell. To address this issue, we determined bacterial cell numbers using fluorescence microscopy and bacterial genome copies using qPCR. We found no differences: Ot IB and EB harbored just over one estimated genome copy per cell (Fig. 1E). This measurement is an approximation that suffers limitations associated with absolute quantification using qPCR and microscopy, but any such errors are expected to be constant between the IB and EB measurements.

Finally, we asked whether IB and EB Ot forms differed in their metabolic activity. Ot is an obligate intracellular bacterium with limited central metabolic and biosynthetic capability^14^ and a reliance on host cells for carbon and nitrogen sources and metabolic intermediates. These observations led us to hypothesize that EB bacteria are metabolically inactive owing to their extracellular location. To test this, we used a microscopy-based assay whereby growing bacteria are exposed to a clickable methionine analog L-Homopropargylglycine (HPG) that is subsequently labeled with a fluorophore^15^ (Fig. 1F). Using this methodology, we observed that 95 % of IB were translationally active five days post-infection, compared with 0% of EB (Fig. 1G). It is possible that EB Ot undergo some residual metabolic activity that is not detected by our assay of protein translation. The fact that not all IB were labelled with HPG indicated that this result did not simply reflect a difference in bacterial permeability between the IB and EB forms. These results provide an additional line of evidence to support the existence of two fundamentally different Ot states.

### IB and EB forms of Ot exhibit different gene and protein expression profiles

We next investigated whether the differences in physical properties and metabolic activity observed in IB and EB forms of Ot were accounted exclusively by physical location relative to host cells. Exposure of EB forms to an osmotically different extracellular environment combined with the lack of nutrient availability might account for our observations. However, a second possibility is that EB forms arise by differentiation of IB into a biochemically distinct state. We reasoned that bacteria in the same developmental stage but differing in position relative to host cells would exhibit almost identical protein profiles. To test this, we performed shotgun mass spectrometric analyses on isolated IB and EB Ot forms. We found that whilst the correlation coefficient of proteins detected in IB and EB isolated four days post-infection was strong (correlation coefficient = 0.843) (Supp. Fig. 3) there were several proteins that accumulated to different levels in the two populations. A complete list of the relative levels of proteins detected in IB and EB is provided in Supp. Table 1. Pathway analyses revealed that IB Ot was enriched in proteins involved in protein synthesis (Fig. 2A). In contrast, EB Ot displayed high expression of SpoT and RpoH, two proteins involved in the stringent response (Fig. 2B). In order to validate these results, we generated antibodies against four proteins that were present at different levels in IB and EB Ot: SpoT, RpoH, RpoD and CtrA. We carried out immunoblot analysis of IB and EB Ot and found that both SpoT and RpoH were present at higher levels in EB than IB, whilst RpoD and CtrA were present at higher levels in IB Ot (Fig. 2C), consistent with the proteomics analysis.

**Fig. 2:**
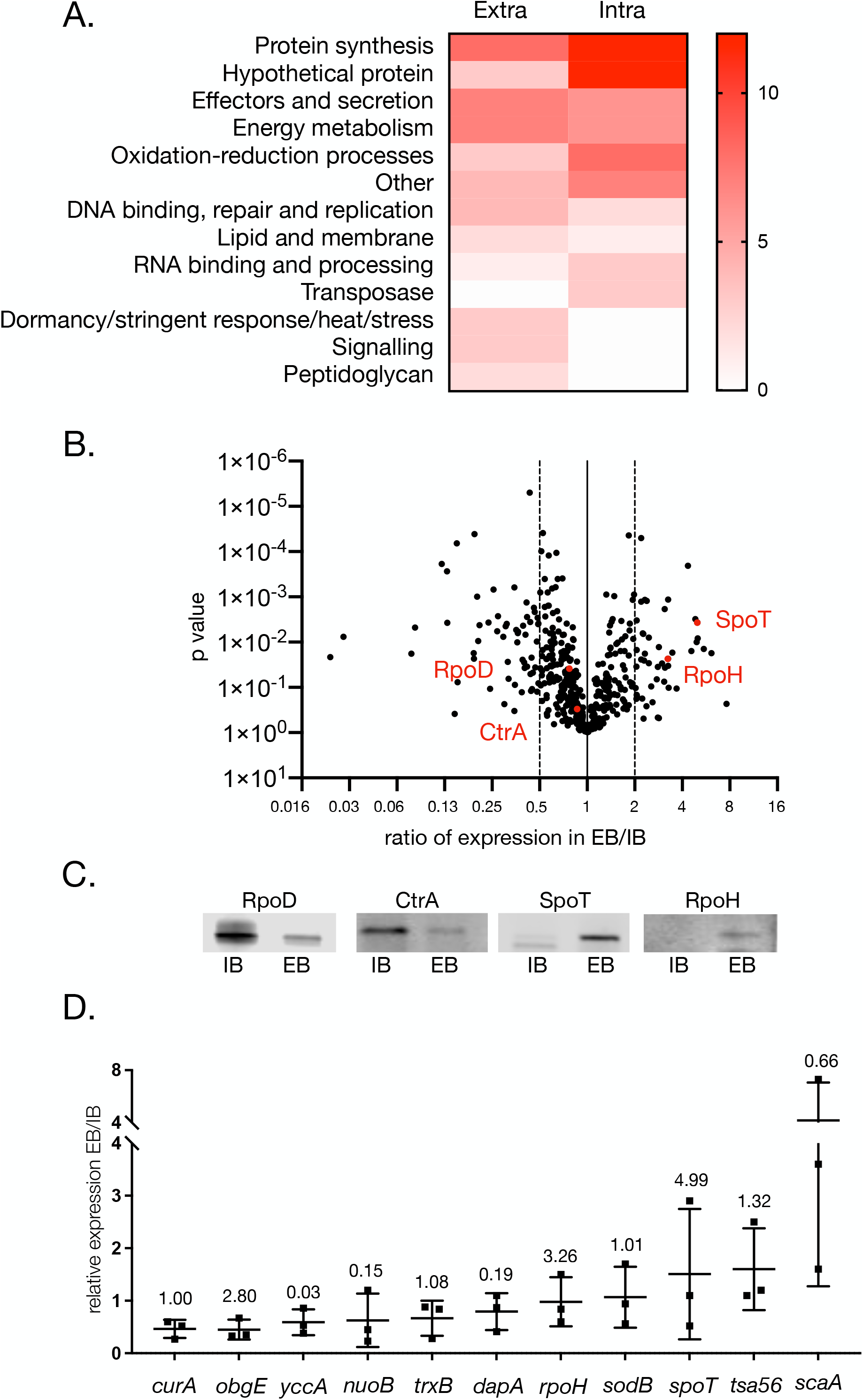
Intracellular and extracellular forms of Ot have different levels of certain proteins. A. Pathway analysis of proteins present at >2X difference between IB and EB forms of Ot (60 proteins in IB, 47 proteins in EB). Bacteria were grown in L929 cells at an MOI of 410:1 and harvested four days post-infection. IB and EB Ot were isolated and analysed by shotgun proteomics. B. SpoT and RpoH are present at higher levels in EB compared with IB Ot as measured by proteomics described in A. C. Immunoblot analysis showing relative levels of four Ot proteins SpoT, RpoD, RpoH and CtrA in IB and EB Ot. Loading was normalised by measuring bacterial genome copy number using qPCR and loading the same number of IB and EB Ot in each pairwise comparison. Samples taken from the same experiment are used for the four antibodies in this figure, with the same amount loaded in each immunoblot. Uncropped immunoblot images showing antibody specificity are given in Supp. Fig. 4. This data confirms the trends observed in the proteomics analysis for these four proteins. D. RTqPCR analysis showing relative levels of 11 genes expressed in EB and IB Ot. Bacteria were isolated from L929 cells four days post-infection at an MOI of 672:1, 300:1, and 560:1. The figure depicts results from three independent replicates. Gene expression levels were normalised using three genes: *mipZ*, *ank12*, and *16S.* Mean and standard deviation values are shown. The ratio of protein levels in EB/IB, as found in the proteomics experiment in 2A, is given above each gene for comparison.

We next investigated whether EB and IB Ot exhibited differences in transcript expression levels. These experiments required the propagation of large amounts of EB because this form of Ot is metabolically inactive and thus harbors very low mRNA levels. We selected 11 genes, three of which (*curA, trxB, sodB*) were expressed at similar levels in our proteomic studies comparing EB and IB. Additionally, we quantified mRNA expression of four genes encoding proteins enriched in IB Ot (*yccA, nuoB, dapA, scaA*) and four genes enriched in EB Ot (*obgE, spoT, tsa56, rpoH*). We isolated RNA from IB and EB bacteria four days post-infection and assessed mRNA levels by qRT-PCR (Fig. 2D). We found no correlation between mRNA and protein levels in multiple cases. For example, *rpoH* transcript levels were similar in IB and EB Ot, but protein levels were higher in EB. These observations provide additional evidence supporting the existence of molecularly distinct bacterial forms and suggest that transcriptional and post-transcriptional regulatory mechanisms participate in the differentiation process.

### Surface proteins are present at different relative levels on IB and EB forms of Ot

Our differential proteomics analyses revealed that IB and EB have different levels of specific surface proteins (Fig. 3A). Surface proteins play a central role in Ot function for two reasons. First, unlike most Gram-negative bacteria, Ot lacks lipopolysaccharide, which generally serves as a membrane shield. Second, Ot is a cytosolic bacterium, and its surface is in direct contact with cytoplasmic host components. Ot harbors genes encoding four widely conserved autotransporter proteins: ScaA, ScaC, ScaD, and ScaE. Using our proteomics dataset, we detected ScaA and ScaC, but not ScaD or ScaE. ScaA was slightly enriched in IB relative to EB Ot (1.5X), while ScaC was enriched in EB (1.87X). Ot also harbors genes encoding three immunogenic surface proteins that dominate the humoral response in human patients, TSA22, TSA47, and TSA56. We found that TSA22 was enriched in IB relative to EB Ot (3.2X) while TSA47 and TSA56 were not present at significantly different levels (1.14X and 1.32X increase in EB compared with IB, respectively). To verify our observations, we generated polyclonal antibodies against ScaA and ScaC and determined their relative levels in IB and EB bacteria using immunoblot analyses (Fig. 3B and Supp. Fig. 4). Our experiments up to this point were performed in mouse fibroblasts (L929) because Ot grows readily in these cells. Moreover, L929 fibroblasts are suitable for characterizing bacterial properties that are not affected by the cellular environment. However, for our validation studies, we also investigated bacteria grown in primary human umbilical vein endothelial cells (HUVEC) as a more physiologically relevant model of Ot infection in humans. Our immunoblot studies in both L929 and HUVECs confirmed enrichment of ScaA in IB Ot and higher levels of ScaC in EB compared to IB Ot, in agreement with our mass spectrometric analyses.

**Fig. 3:**
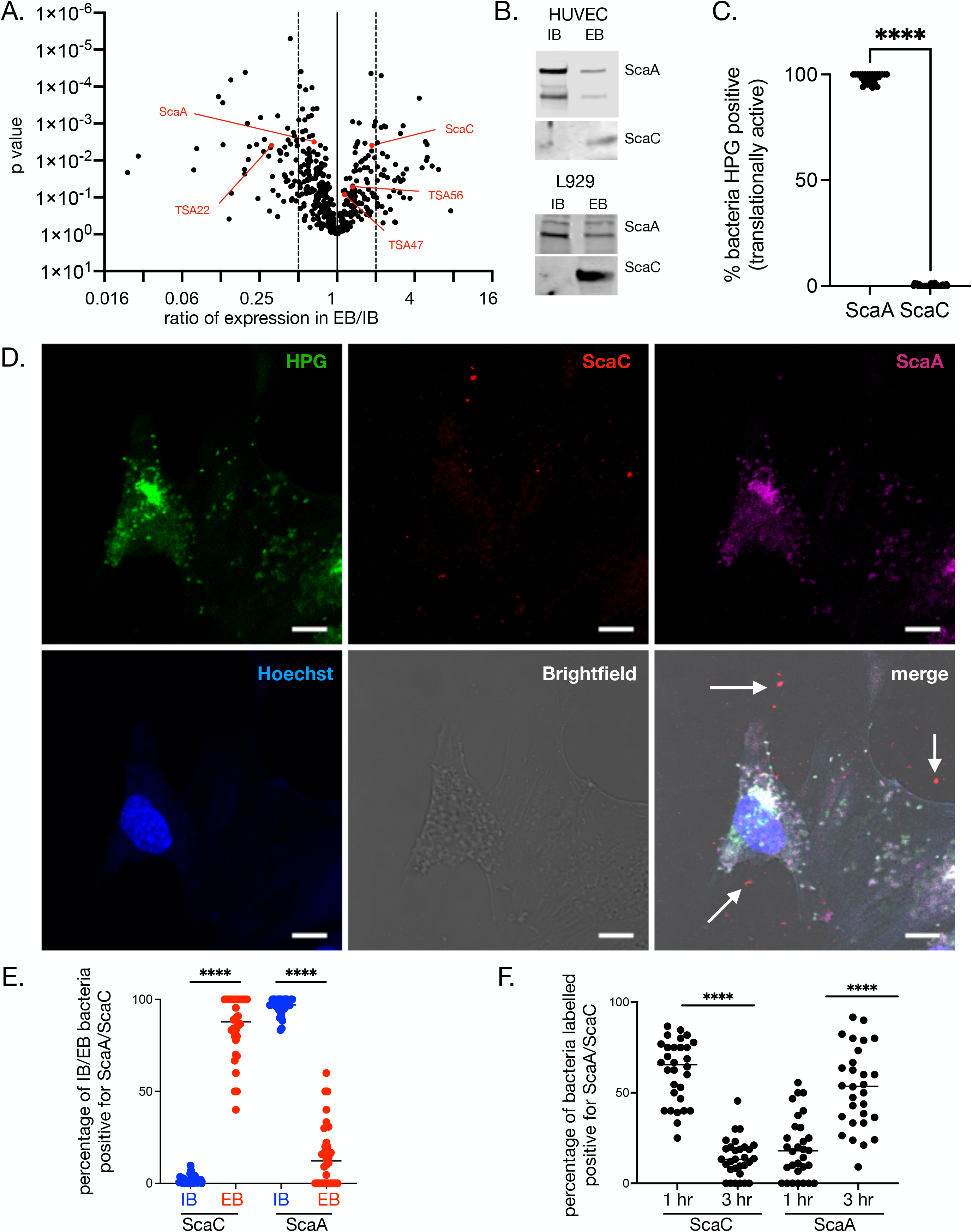
Autotransporter proteins ScaA and ScaC are differentially expressed on IB and EB forms of Ot. A. Volcano plot showing differential expression of surface proteins in proteomics analysis of IB and EB forms of Ot. We observed higher expression of ScaC, TSA56, and TSA47 in EB whilst IB Ot expressed higher ScaA and TSA22 levels B. Immunoblot analysis of IB and EB Ot from infected L929 and HUVEC cells confirming differential expression of ScaA, and ScaC between IB and EB forms. Loading was normalised by measuring bacterial genome copy number using qPCR and loading the same number of IB and EB Ot in each pairwise comparison. Uncropped immunoblot images showing antibody specificity are given in Supp. Fig. 4. C. Quantification of translational activity, as determined by a positive confocal microscopy signal observed with the clickable methionine probe L-homopropylgylglycine (HPG). HUVEC cells were infected with Ot for five days, then fixed and labelled with HPG and anti-ScaA or anti-ScaC antibodies. ScaA/ScaC-positive bacteria were scored as being positive or negative for HPG labelling. D. Confocal microscopy images of HUVEC cells infected with Ot for five days and labelled for DNA (Hoechst, blue), translational activity (HPG, green), ScaA (anti-ScaA antibody, magenta) and ScaC (anti-ScaC antibody, red). Scale bar = 10 μm. Examples of extracellular bacteria labelled with ScaC are shown with a white arrow. More images as well as equivalent experiments in L929 cells are shown in Supplementary Figures 5-7. E. Quantification of confocal micrographs of HUVEC cells infected with Ot and labelled as in 3D. The percentage of IB and EB Ot at five days post-infection that were positively labelled with anti-ScaA and -ScaC antibodies was counted. F. Quantification of ScaA- and ScaC-positive IB bacteria inside HUVEC cells one- and three-hours post-infection, showing an increase in ScaA labelling and a decrease in ScaC labelling. Throughout this figure quantification was carried out by randomly imaging 30 fields across three independent experiments. Graphs show individual values and means. Statistical significance was calculated using an unpaired t test with Graphpad Prism software. * p-value ≤ 0.05, ** p-value ≤ 0.01, *** p-value ≤ 0.001, **** p-value ≤ 0.0001.

### ScaA and ScaC as markers of EB and IB forms of Ot

Our observation that 95% of IB is translationally active five days post-infection, compared with 0% of EB, combined with the fact that these Ot forms display opposite relative levels of ScaA and ScaC led us to investigate whether these surface proteins can be used as *bonafide* markers of metabolically active IB and metabolically quiescent EB Ot forms, respectively. We used immunofluorescence microscopy to identify bacteria expressing ScaA and ScaC and then quantified the number of translationally active ScaA- and ScaC-positive bacteria by measuring incorporation of HPG. We found (Fig. 3C and Supp. Fig. 7A) that ScaA-positive bacteria, in contrast to ScaC-positive Ot, were mostly translationally active. We then quantified the number of ScaA/ScaC positive bacteria inside and outside cells and confirmed that ScaC and ScaA levels were higher in EB and IB Ot, respectively (Fig. 3D, E and Supp. Fig. 5, 6, 7B). Together, these results show that ScaA and ScaC are present at different relative levels depending on bacterial location relative to host cells: metabolically active IB exhibit higher levels of ScaA relative to EB Ot. Conversely, levels of ScaC in metabolically inactive EB is high relative to that of IB forms of Ot.

The observation that EB exhibits high *scaA* transcript levels (Fig. 2D) but low levels of ScaA protein led us to hypothesize that ScaA is translated into protein soon after entry into host cells, using pre-formed mRNA. We quantified the number of IB Ot expressing ScaA and ScaC at 1- and 3-hours post-infection and found a decrease in ScaC levels and an increase in ScaA levels as the infection proceeded (Fig. 3F and Supp. Fig. 7C). These results show that ScaA accumulates during the first 3 hours after infection, suggesting that *scaA* transcripts are made in EBs in anticipation of new infections. These results point to the existence of regulatory mechanisms that control levels of ScaA and ScaC on the surface of Ot cells, depending on bacterial location and stage of host infection.

### IB and EB Ot populations are infectious but differ in physical stability and mechanisms of entry into host cells

Previous studies have shown that specific Anaplasmataceae differentiate into distinct intracellular and extracellular forms and that only the extracellular form is effective at infecting host cells^16^. This observation led us to investigate whether the ability to infect host cells differs in IB compared with EB Ot. Growth curve measurements revealed that both bacterial forms could infect L929 and HUVEC cells and replicate with similar dynamics (Fig. 4A and Supp. Fig. 8). However, IB appears to retain infectivity more efficiently than EB following physical and osmotic challenges. Specifically, when we subjected IB and EB Ot to physical stress using a bead mill for 1-2 minutes, we found that EB displayed a reduction in growth that was dependent on the time of exposure to stress; in contrast, IB was unaffected by this treatment (Fig. 4B). Furthermore, osmotic shock resulted in a significantly greater reduction in viability in EB compared with IB (Fig. 4C). Together, these results demonstrate that IB is physically more robust than EB; this property is likely due to differences in peptidoglycan content, as described above.

**Fig. 4:**
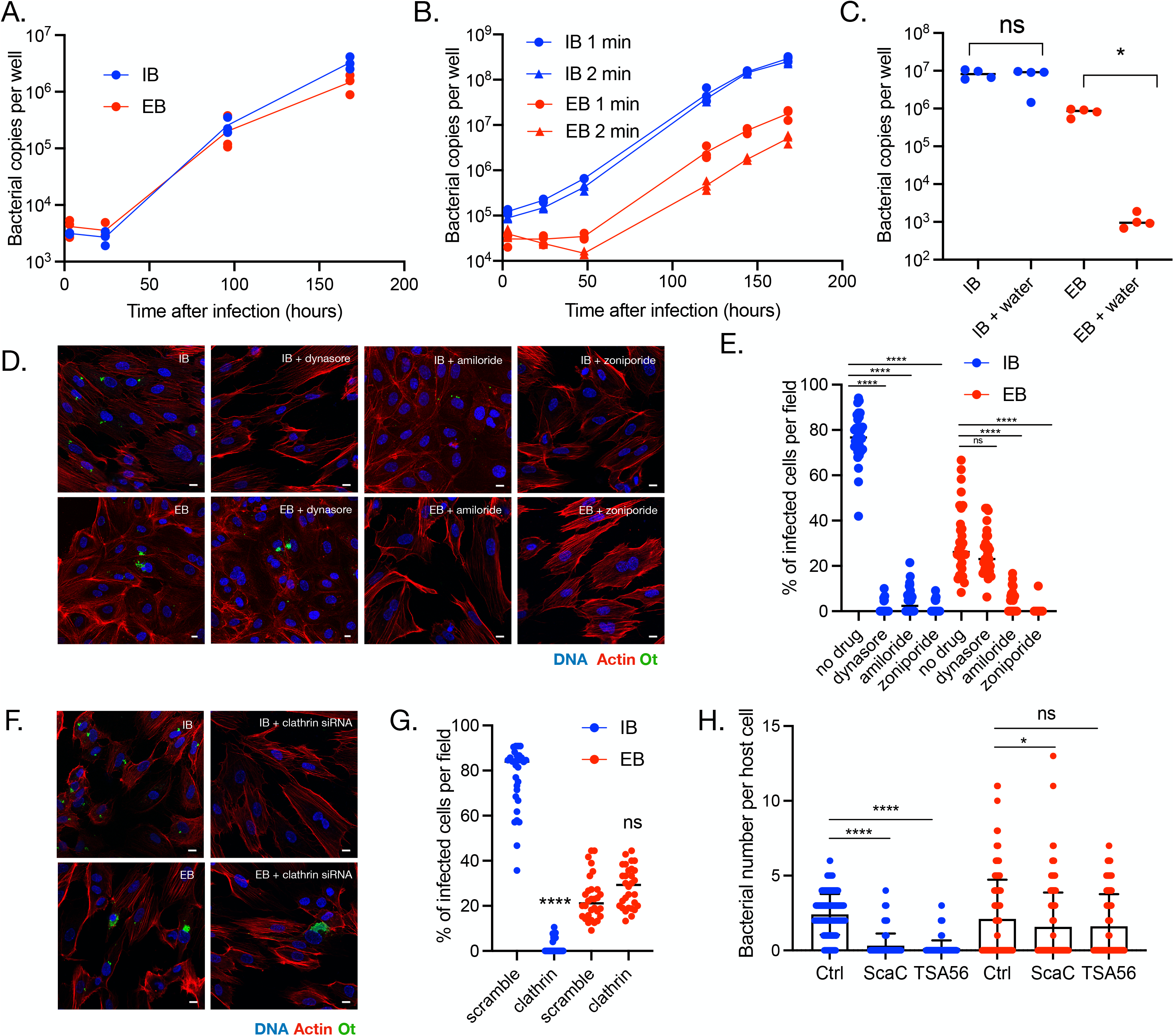
Both IB and EB populations are infectious but differ in physical stability and mechanism of entry into cells. IB and EB Ot were isolated from infected L929 cells five days post-infection and grown in fresh L929 cells in a 24-well plate. The number of intracellular bacteria was then quantified at 3-, 24-, 96-, and 168-hours post-infection using qPCR against *tsa47,* a single copy Ot gene. Doubling times of IB and EB Ot were similar. Similar results were found in HUVEC cells (Supp. Fig. 8). B. IB and EB were exposed to physical stress using a bead mill for one or two min; growth was quantified as in 4A. The bead mill did not affect the growth rate of IB Ot but reduced that of EB Ot in an exposure-dependent manner. C. IB and EB were exposed to osmotic shock by suspending the bacteria in water for five min and then growing them in fresh L929 cells. Bacterial copy number after seven days was quantified by qPCR. EB was more susceptible to osmotic shock than IB Ot, as shown by a more significant decrease in bacterial number. Statistical significance was determined using a Mann-Whitney test. For panels A-C each datapoint was taken from a separate well from a 24-well plate. Images (D.) and quantification (E.) of the effect of endocytosis inhibitors on entry of IB and EB Ot into HUVEC cells. IB and EB were isolated from L929 cells five days post-infection and then seeded onto fresh HUVEC cells that had been treated with inhibitors of clathrin-mediated endocytosis (100 µM dynasore) or macropinocytosis (50 µM amiloride or 100 µM zoniporide). Cells were fixed three hours post-infection. Dynasore inhibits entry of IB but not EB, indicating that clathrin-mediated endocytosis is the major mode of entry of this population, whilst amiloride and zoniporide inhibited entry of both IB and EB. This observation points to a role for macropinocytosis, but not clathrin-mediated endocytosis, in EB’s entry into mammalian cells. Red = actin (phalloidin), blue = Hoechst (DNA), green = Ot (anti-TSA56 antibody). Scale bar = 10 µm. Similar results were found in L929 cells (Supp. Fig. 10). Images (F) and quantification (G) of the effect of 0.2 µM siRNA against clathrin on entry of EB and IB Ot into HUVEC cells. Experiments were carried out as in D. and E. Quantification of E. and G. were carried out as follows: 30 fields were randomly imaged over 3 independent experiments (10 fields per experiment). In each field the percentage of infected host cells was counted. Individual values and means are shown here. Statistical significance in (E) was determined by comparing all groups using one-way ANOVA, followed by a Dunnett’s multiple comparison tests with the control within the IB or EB group, and in (G.) by performing a Mann Whitney test comparing the scramble control and the clathrin siRNA treated condition for IB and EB separately, using GraphPad Prism software. H. Blocking the bacterial surface adhesins TSA56 and ScaC with anti-TSA56 and anti-ScaC antibodies strongly inhibits entry of IB but not EB Ot in HUVEC cells. Bacterial entry in the presence of vehicle control or antibodies was determined by quantifying the number of intracellular bacteria per host cell 30 minutes post infection using immunofluorescence microscopy of bacteria labelled with HPG. Statistical significance was determined by comparing all groups using one-way ANOVA, followed by a Dunnett’s multiple comparison tests with the control within the IB or EB group. **** p ≤ 0.0001; ns = not significant. * p-value ≤ 0.05, ** p-value ≤ 0.01, *** p-value ≤ 0.001, **** p-value ≤ 0.0001. Throughout figure blue = IB and red = EB.

To investigate whether EB and IB forms of Ot enter host cells using the same invasion mechanism, we pre-treated HUVEC cells with the clathrin-mediated endocytosis inhibitor dynasore or the macropinocytosis inhibitors amiloride and zoniporide. We then infected the cells with IB or EB Ot and quantified the number of intracellular bacteria 3 hours post-infection using confocal microscopy (Fig. 4D, E). The entry of IB Ot into the cells was blocked by dynasore, thus confirming the participation of clathrin-mediated endocytosis in this process. IB entry was also inhibited by amiloride and zoniporide, suggesting additional involvement of macropinocytosis in bacterial uptake. In contrast, EB Ot entry into cells was not affected by dynasore treatment but was completely inhibited by pre-incubation with amiloride or zoniporide. To confirm that EB Ot did not use clathrin-mediated endocytosis to enter cells, we treated HUVEC and L929 cells with siRNA against clathrin. Control microscopy experiments demonstrated that clathrin-targeting siRNA reduced the levels of clathrin foci in uninfected cells (Supp. Fig. 9). We found that entry of IB form Ot required clathrin, whilst entry of EB form Ot did not (Fig. 4F, G and Supp. Fig. 10). Together these results demonstrate that IB form Ot uses clathrin-mediated endocytosis to enter cells whilst EB form Ot does not.

To further demonstrate that IB and EB Ot were entering host cells using distinct mechanisms, we examined the role of the surface adhesins TSA56 and ScaC on bacterial entry. These are outer membrane proteins that have been shown to be involved in entry of Ot into host cells, in experiments carried out using what we now describe as IB Ot stage bacteria^6,7^. We used antibodies targeting ScaC and TSA56 and determined whether these antibodies would block the entry of Ot into host cells. Our results with IB Ot replicated published studies and showed that blocking ScaC and TSA56 using antibodies led to a strong reduction in bacterial entry (Fig. 4H). With EB Ot, however, only a minimal effect was observed (Fig. 4H). This result supports a model in which IB Ot enters host cells using adhesin-induced clathrin-mediated endocytosis, whilst in EB Ot the surface proteins are shielded by host derived membrane and use a different mechanism of entry.

## Discussion

Our studies have unveiled a new feature of the obligate intracellular bacterium Ot. This organism’s unique exit strategy, which involves budding off the surface of infected cells, is associated with differentiation into a molecularly and functionally distinct developmental state (Fig. 5). While unusual in the family Rickettsiaceae, this property is reminiscent of the biphasic lifecycle of multiple species in the sister family Anaplasmataceae^4^ and the unrelated obligate intracellular bacteria *Chlamydia* spp.^17^. However, in contrast to these organisms, Ot resides in the host cytosol in naked form rather than in a membrane-bound vacuole. To our knowledge, this is the first description of an intracellular bacterium living freely in the host’s cytosol and having a biphasic lifecycle. Our findings have important implications for our understanding of Ot biology and unveil opportunities to develop better therapeutic strategies to treat scrub typhus.

**Fig.5:**
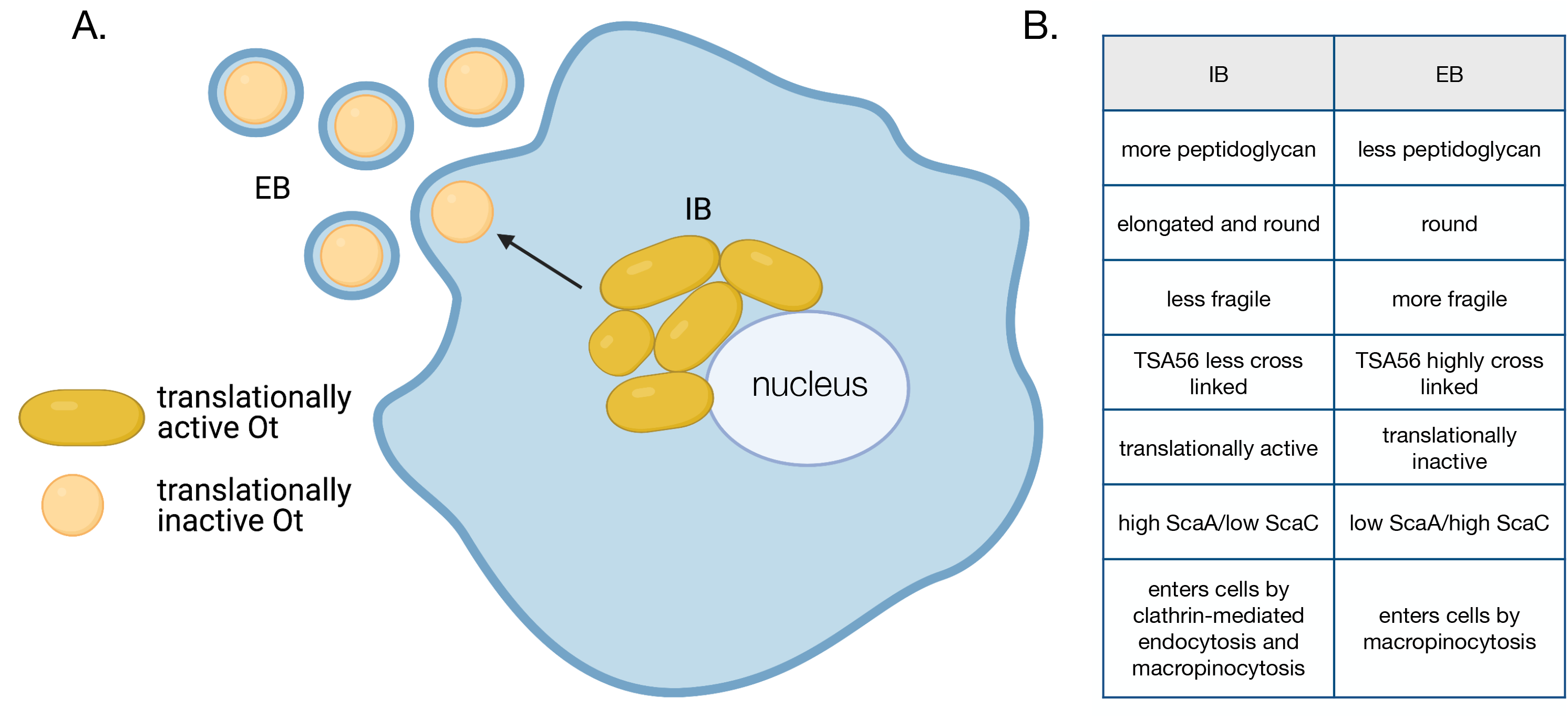
An overview of the infection cycle of Ot, showing differentiation between IB and EB forms. IB replicate in a micro-colony adjacent to the nucleus where they are elongated in shape and translationally active. Bacteria move to the cell’s edge and transition to an EB form that is round and translationally inactive. Created with BioRender.com B. Table summarizing the differences between IB and EB forms of Ot.

The physical properties of prokaryotic cell walls are crucial for survival. These stiff structures maintain cellular shape, protect the interior, counteract osmotic pressure, and provide structural strength through peptidoglycan layers. Our studies revealed that EB and IB forms of Ot differ in peptidoglycan content. Lower levels in EB compared with IB make this form more susceptible to physical and osmotic stress. There is a long-standing question regarding the role of peptidoglycan in obligate intracellular bacteria. These organisms reside in an osmotically protective niche and are under selective pressure to reduce the abundance of immunogenic pathogen-associated molecular patterns (PAMP)s^18^. In the case of pathogenic *Chlamydia* species, it has been shown that peptidoglycan is retained only at the septum and is required for cell division^19^. Higher peptidoglycan levels in IB Ot *vs.* EB Ot point to the retention of peptidoglycan for growth and division rather than structural integrity maintenance.

The observed differences in physical properties between EB and IB forms are complemented by measurable differences in the accumulation of certain proteins, suggesting that each form performs unique functions or responds to specific challenges. Proteomics analyses in EB Ot revealed increased levels of heat shock sigma factor RpoH and the alarmone (p)ppGpp synthetase SpoT associated with the stringent response^20^. These results suggest that the EB form of Ot shares features with the dormant state of other bacteria. Notably, Ot is the only reported member of the Rickettsiaceae to encode a full-length *spoT* gene^4^. This exclusive feature is in harmony with the fact that developmental differentiation of Ot is highly unusual in members of this family. The changes associated with differentiation into the EB state likely serve one or both of the following purposes: ensuring survival in the extracellular state and priming the bacteria for reinfection of new cells. Indeed, the observation that *scaA* transcripts are abundant in the EB form and that ScaA protein is synthesized early upon entry into host cells supports the idea that EB Ot is primed for infection.

Comparing the protein and RNA transcript levels of selected genes revealed a poor correlation for many genes. A potential explanation is that RNA levels are low in the metabolically inert EB form, which does not reflect newly translated proteins. Ot strain UT76 encodes only 8 predicted transcription factors and two sigma factors, RpoD and RpoH^21^. Thus, it is possible that gene expression is not strictly regulated at the transcriptional level but that other forms of regulation are prevalent in Ot. We recently showed high antisense transcription levels in Ot, especially in genes associated with the rampant integrative conjugative element RAGE, which dominates the Ot genome^22^. There, we found that high antisense transcription correlates with low protein levels for a particular gene, pointing to this mechanism as a central regulatory strategy in Ot. Gene expression in the Ot-related, free-living alphaproteobacterium *Caulobacter crescentus* is highly regulated by proteolysis^23^. Mechanisms based on differential proteolysis may therefore also play an essential role as critical determinants of protein levels in Ot.

Our observation that EB Ot enters cells through pinocytosis, whereas IB Ot utilizes endocytosis and, to a lesser extent, pinocytosis resolves a long-standing gap in the field as it was unclear how budded bacteria enveloped in plasma membrane enter host cells. Physiologically, IB entry into host cells may be necessary during trafficking into adjacent cells, as previously described for spotted fever group *Rickettsia* spp.^24^. Additionally, low levels of cell lysis may release IB Ot forms that can then infect new host cells. The different mechanisms utilized by EB and IB to enter cells suggest that strategies to escape from the endolysosomal pathway will differ between the two Ot forms. It is unclear whether and how the differing transcriptional and proteomic profiles at the point of infection will affect early host-pathogen interactions. Our finding that IB and EB’s growth dynamics are similar suggests that early differences do not persist through the infection cycle.

In summary, our finding that Ot differentiates into molecularly and functionally distinct forms provides a novel framework to study the infection cycle of this important obligate intracellular pathogen and opens avenues to investigate the physiological significance of the distinct populations characterized.

## Methods

### Cell culture, bacterial propagation and quantification, and reagents

Proteomics and qRT-PCR experiments were performed using the mouse fibroblast cell line L929 kindly provided by Dr. Blacksell at the Mahidol Oxford Tropical Medicine Research Unit, Bangkok, Thailand. For all other experiments, we used L929 cells (ATCC CCL-1), Primary Human Umbilical Vein Endothelial Cells (HUVEC) (ATCC PCS-100-010), monkey kidney epithelial Vero cells (ATCC CRL-1586) or human dermal microvascular endothelium HMEC-1 cells (ATCC CRL-3243), all purchased from ATCC. We cultured cells in either DMEM (L929) or Large Vessel Endothelial Supplement (LVES, HUVEC) supplemented with 10% FBS (Gibco 16140071) at 37°C and a 5% CO_2_ atmosphere. The Karp-like clinical isolate strain of *Orientia tsutsugamushi* UT76^21^ was used throughout the study. For routine propagation, bacteria were grown in 25- or 75-cm^2^ culture flasks, as described previously^12^. We extracted DNA using alkaline lysis buffer^12^, and then assessed bacterial quantities by determining genome copy numbers relative to a standard curve. We used a primer/probe set to amplify the single copy gene *tsa47* (tsa47F 5’-TCCAGAATTAAATGAGAATTTAGGAC-3’, tsa47R 5’-TTAGTAATTACATCTCCAGGAGCAA-3’, tsa47probe 5’-[6FAM]TTCCACATTGTGCTGCAGATCCTTC[TAM]-3’) or the 16S gene (Rick16SF 5’-5’-GCTACACGCGTGCTACAATGG-3’, Rick16SR 5’-TGTGTACAAGGCCCGAGAACG-3’, Rick16Sprobe 5’-[6FAM] ATCGCTAGTAATCGCGGATCAGCATGCC [TAM]-3’). For drug inhibition assays, dynasore (Sigma D7693) and amiloride (Sigma A3085) were added to L929 or HUVEC cells 1 hour before infection with Ot.

### Fluorescence microscopy

#### Immunofluorescence labelling and confocal microscopy

Following fixation, cells were permeabilized by incubation in 0.5% Triton X-100 for 30 min on ice followed by blocking in 1XPBS (pH7.4) containing either 1 mg/ml BSA or sera (diluted 1:20) matching the labeled antibody’s host, for 30 min at room temperature. Incubations with primary antibodies took place for 1 hour at 37°C. These studies were conducted using in-house-generated antibodies (*i.e.*, rat monoclonal antibody against TSA56 diluted 1:200; rabbit polyclonal antibody against Ot ScaC diluted 1:200) or commercial products (*i.e.*, MA1-164, a Thermo Scientific monoclonal mouse antibody against LAMP1 diluted 1:200). The fixed cells were washed twice with PBS-BSA and were then incubated with appropriate secondary antibodies (1:1000 dilutions) for 30 min at 37°C in the dark. We used the following secondary antibodies: goat anti-mouse IgG superclonal Alexa Fluor 647 conjugate (Thermo Fisher A28180), goat anti-rat Alexa Fluor 488 (Thermo Fisher A11006), and goat anti-rabbit Alexa Fluor 594 (Thermo Fisher A11012). The nuclear stain Hoechst (1:1000 dilution) was supplemented to incubations with secondary antibodies. Phalloidin Alexa Fluor 594 (1:200, ThermoFisher Scientific A12381) was used to detect host actin. The cells were washed with 1X PBS before supplementation of mounting medium [20 mM Tris (pH 8.0); 0.5% N-propyl-gallate; 90% glycerol]. Imaging was performed using a Zeiss Observer Z1 LSM700 confocal microscope with an HBO 100 illuminating system equipped with a 63x/1.4 Plan-APOCHROMAT objective lens (Carl Zeiss, Germany) and 405 nm, 488 nm, and 555 nm laser lines. In some cases, we used a Leica TCS SP8 confocal microscope (Leica Microsystems, Germany) equipped with a 63x/1.4 Plan-APOCHROMAT oil objective lens with a 1.4-mm working distance and 405 nm, 488 nm, 552 nm, and 638 nm laser lines.

Fluorescence intensity measurements were performed manually using Fiji ImageJ software. 30 (Fig. 3C) or 100 (Fig. 1G); individual bacteria were quantified by measuring the brightest pixel intensity within the bacterial area. Images from individual experiments were acquired under identical laser intensity and acquisition conditions. Length and width measurements were performed manually using Fiji Image J software.

### STORM imaging and analysis

We acquired 3D-STORM data using a homebuilt multi-color Total Internal Reflection Fluorescence (TIRF) microscope^25^ that is based on an inverted microscope (Nikon, Ti-E), a high NA objective (Nikon, CFI-Apo 100X, NA 1.49), a 405 nm photoactivation laser (Coherent, OBIS 405 LX 100mW), a 488 nm excitation laser (MPB, 2RU-VFL-P-20000642-B1R), and an EMCCD camera (Andor, iXon Ultra-888). Samples immunolabelled with Alexa 488 were mounted on the microscope, transferred to STORM buffer [OxyFluor (Oxyrase), 50 mM betamercaptoethanol (Sigma), 2.5 mM cyclooctatetraene (COT) (Sigma)], and illuminated with a strong (488 nm) laser to record single-molecule fluorescence blinking events at a frame rate of 25 Hz. Illumination at 405 nm was intercalated with pulsed 488nm excitation to enhance photoactivation of Alexa 488 without exacerbating background signal. A cylindrical lens (Thorlabs, LJ1516RM-A) was placed in front of the camera to determine the z-coordinates of fluorescent molecules through the elongated point spread function (PSF) methods^26^. Each STORM image was reconstructed from single-molecule localization data obtained by analysing 30,000 frames with custom Matlab codes. Briefly, bright single fluorescence spots were identified by thresholding methods, and their 3-D coordinates were determined by 2-D elliptical Gaussian fitting along with a pre-calibrated conversion table between eccentricity and z-coordinate. Focus drift was actively stabilized with Perfect Focus System (Nikon) while acquiring the data, whereas xy-drift was post-corrected using a redundant cross-correlation algorithm^27^. Each localization event was represented as a 2-D Gaussian of 20 nm standard deviation in STORM images. For the analysis of bacterial morphology, we selected cells that showed clear cell boundary, manually adjusted the z-coordinate to locate the cell midplane, and measured long- and short-axes length.

#### Click labelling

Our metabolic click-labelling was based on the Click-iT HPG Alexa Fluor protein Synthesis Assay Kit (Click-iT HPG Alexa Fluor Protein Synthesis Assay Kit, ThermoFisher C10428) strategy. To incorporate L-homopropargylglycine (HPG), we incubated infected cells with minimal medium (Dulbecco’s Modified Eagle Medium, DMEM, ThermoFisher 21013) lacking L-methionine and containing 25 µM HPG, for 30 min at 37°C. Labelled bacteria were washed twice with PBS containing 1mg/ml BSA, fixed with 1% formaldehyde, and subsequently permeabilized with 0.5% Triton X-100 for 20 min on ice. After further washing with PBS containing 1 mg/ml BSA, we supplemented the Click-iT reaction cocktail (Click-iT HPG Alexa Fluor Protein Synthesis Assay Kit, ThermoFisher C10428) and incubated the cells for 30 min at room temperature in a light protected environment. The Azide dye (Alexa Fluor 488) was used at a final concentration of 5 µM. Following completion of the click reaction, the cells were subjected to immunofluorescence labelling and imaging as above.

### Gene expression analysis by RT-qPCR

L929 cells were infected with *Orientia tsutsugamushi* UT76 using multiple MOIs to a target MOI of 400:1 (quantitative PCR on freshly isolated Ot showed variation of actual MOIs ranging from 200:1 to 672:1 across 3 independent infections). Very high MOIs were used due to the low levels of EB bacteria combined with the fact that RNA levels in EB Ot were very low. Infected cells were harvested four days post infection and were stored in RNAProtect Bacteria Reagent (Qiagen 76506) at -80°C. Total RNA was extracted using the Qiagen RNeasy Plus kit (Qiagen 74136) according to the manufacturer’s instructions. One μg of total RNA was incubated with DNaseI (Thermo Fisher Scientific, AM2238) for 16 h at 37°C, and the samples were then reverse-transcribed using the iScript reverse transcription supermix (Biorad 170–8841) and random primers. Controls lacking reverse transcriptase demonstrated that the RNA sample used for cDNA synthesis was not contaminated with genomic DNA. The resulting cDNA was then used as template for qPCR using primers for *spoT, rpoH, obgE, yccA, nuoB, dapA, curA, sodB, trxB, tsa56, and scaA.* RT-qPCR was performed using SYBR green qPCR mix (Biotools 10.609). Expression levels were normalized relative to the housekeeping genes *16sRNA, ank12_1 and mipZ*.

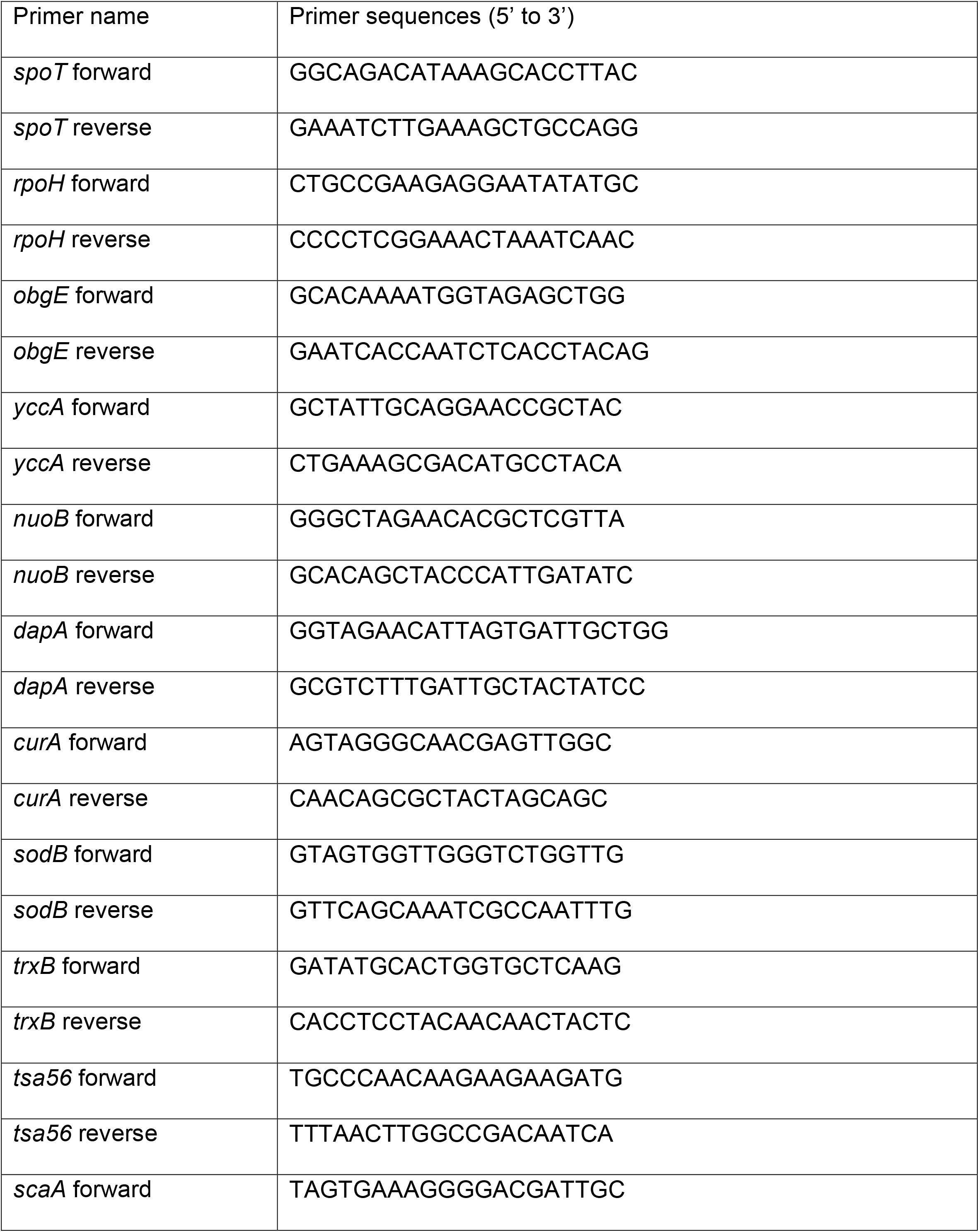
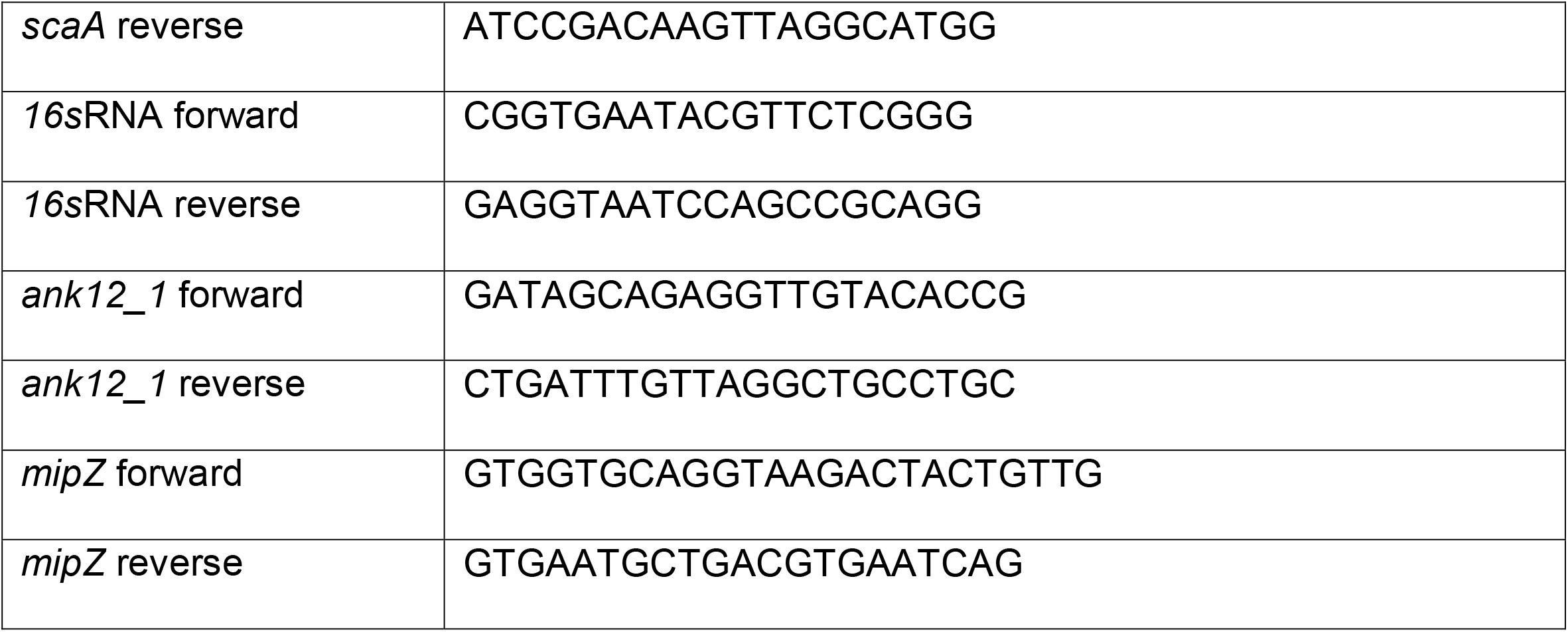

### Proteomics sample preparation

We used *Orientia tsutsugamushi* strain UT76 to infect L929 mouse fibroblasts at a 410:1 MOI in six independent culture flasks. High MOIs were used to ensure sufficient material for proteomics analysis. Despite the high MOI the L929 cells remained intact and attached to the surface of the flask at the point of harvesting the samples. We replaced the culture medium at one day postinfection (dpi) to remove bacteria that failed to enter the host. Intracellular and extracellular bacteria were isolated at four dpi; two separate flasks were harvested as one replicate, yielding a total of three replicates. L929 cells were grown in 75-cm^2^ plastic flasks at 37°C in a 5% CO_2_ atmosphere, using DMEM media (Gibco 11965092) supplemented with 10% FBS (Sigma-Aldrich F7524). At four dpi, the media were transferred to 50-ml conical tubes for extracellular bacteria isolation. Culture flasks were rinsed with 15 ml of PBS, and cells were scraped using two ml of fresh media per flask. The detached cells were transferred to two-ml Safe-lock tubes (Eppendorf 0030120094) and homogenized for three minutes using a Bullet blender^®^ homogenizer (Next Advance #BBY24M) with speed control setting at 8. Host cell debris was removed by centrifugation at 300 g for three minutes; we then harvested intracellular bacteria (IB) by an additional three-minute centrifugation at 20,000 g. Before isolation of extracellular bacteria (EB), we removed possible detached host cells containing IB through a pre-spin step (1,000 g, 5 min, 4°C). No visible pellet was observed indicating that there were negligible amounts of detached host cells potentially contaminating the EB prep with IB bacteria. Following transfer of supernatants to new conical tubes, we harvested extracellular bacteria by centrifugation at 3000 g for 20 minutes at 4°C. IBs and EBs were washed with 0.3 M sucrose three times, resuspended in PBS, and lysed with 1% Triton X-100 for 30 minutes at 4°C. Bacterial proteins were precipitated by addition of 80% ice-cold acetone and overnight incubation at -20°C. Protein pellets were washed in acetone twice to remove residual Triton X-100 by centrifugation at 15,000 g for 10 min. Briefly dried protein extracts were then dissolved in 100 µl of 50% trifluoroethanol in 100 mM Triethylammonium bicarbonate (TEAB) buffer. The samples were reduced by treatment with 20 mM tris(2-carboxyethyl)phosphine (TCEP) for 20 minutes at 55°C, alkylated in 55 mM chloroacetamide (CAA) at room temperature in the dark, and digested with endoproteinase LysC (LysC: protein ratio=1:50) overnight at 37°C and, subsequently, with trypsin (trypsin: protein ratio=1:100), also overnight at 37°C. Average bacterial copy numbers for IBs and EBs were estimated by quantitative PCR yielding 1.5E+08 and 2.6E+09 per replicate, respectively. The whole protein content in each replicate was used without normalisation at this step due to the fact that IBs and EBs were isolated from different environment and therefore have diverse amounts of co-purified host protein. Samples were acidified with Trifluoroacetic acid (TFA) to 1% final concentration and centrifuged to remove insoluble materials. Trypsinized samples were desalted using an Oasis^®^ HLB 1cc cartridge (Waters™). Briefly, following cartridge activation with 500 µl of 100% acetonitrile and equilibration with 500 µl of 0.5% acetic acid, we loaded samples and then washed the cartridge with 1 ml MS-grade water. Peptides were eluted with 500 µl of 80% acetonitrile in 0.5% acetic acid, then transferred to low-binding microtubes, and dried using a SpeedVac. Each peptide pellet was resuspended with 20 µl of MS-grade water. We estimated peptide content using 5% of each sample, so called ‘Mini-runs’. This was to check whether we need to adjust or normalise sample loadings in the actual runs.. Briefly, individual aliquots in 1 µl of peptides were supplemented with 4 µl of 1% (v/v) acetonitrile containing 0.06% (v/v) trifluoroacetic acid and 0.5% (v/v) acetic acid, transferred to a 96-well autosampler plate, and then injected (3 µl) into an Orbitrap Fusion™ Mass Spectrometer. After we screened out host proteins from the Mini-runs result, the frequency distribution of Ot protein abundances were plotted. Medians of IB and EB replicates showed abundance values with maximum difference of 1 log scale, suggesting a single normalisation process may be performed during data analysis. We decided to fractionate the rest of samples without the need of normalizing sample loadings. The remaining 95% of each sample (19 µl in MS-grade water) was mixed with 20 µl of 10 mM ammonium formate (pH 10) and centrifuged at 20,000 g for 3 min to eliminate insoluble components. Each peptide sample was then fractionated through a step-wise desalting process on C18 stage tips using 10%, 15%, 25%, and 50% acetonitrile in 10 mM ammonium formate (pH 10). Fractionated samples were washed with 60% acetonitrile containing 0.1% formic acid twice to eliminate residual ammonium formate by evaporation before tandem MS analysis.

### LC-MS/MS

Dried samples were resuspended in 10 µl of 2% (v/v) acetonitrile containing 0.06% (v/v) trifluoroacetic acid and 0.5% (v/v) acetic acid, and 1 µl was loaded onto an autosampler plate. Online chromatography was performed in an EASY-nLC 1000 (ThermoScientific) instrument using a one-column setup and 0.1% formic acid in water / 0.1% formic acid in acetonitrile as mobile phases. The samples were injected and separated on a reversed-phase C18 analytical column (EASY-Spray LC Column, 75 µm inner diameter × 50 cm, 2 µm particle size; ThermoScientific) maintained at 50°C and using a 2-23% (v/v) acetonitrile gradient over 60 min, followed by an increase to 50% over the next 20 min, and to 90% over 5 min. The final mixture was maintained on the column for 5 min to elute all remaining peptides. Total run duration for each sample was 90 min at a constant flow rate of 300 nl/min.

Data were acquired using an Orbitrap Fusion Lumos™ mass spectrometer (ThermoScientific) using the instrument’s data-dependent mode. Samples were ionized using 2.5 kV and 300°C at the nanospray source; positively-charged precursor MS1 signals were detected using an Orbitrap analyser set to 60,000 resolution, automatic gain control (AGC) target of 400,000 ions, and maximum injection time (IT) of 50 ms. Precursors with charges 2-7 and having the highest ion counts in each MS1 scan were further fragmented using collision-induced dissociation (CID) at 35% normalized collision energy. MS2 signals were analysed by ion trap at an AGC of 15,000 and maximum IT of 50 ms. Precursors used for MS2 scans were excluded for 90 s to avoid re-sampling of high abundance peptides. The MS1–MS2 cycles were repeated every 3 s until completion of the run.

### Proteomic data analysis

Proteins were identified using Proteome Discoverer™ (v2.3, Thermo Scientific). Raw mass spectra were searched against *Orientia tsutsugamushi* primary protein sequences derived from complete genome data for the UT76 strain. Mouse whole proteome sequences were obtained from Uniprot and included as background. Carbamidomethylation on Cys was set as a fixed modification; deamidation of asparagine and glutamine, acetylation on protein N termini, and methionine oxidation were set as dynamic modifications for the search. Trypsin/P was set as the digestion enzyme and was allowed up to three missed cleavage sites. Precursors and fragments were accepted if they had a mass error within 10 ppm and 0.8 Da, respectively. Peptides were matched to spectra at a false discovery rate (FDR) of 1% against the decoy database.

For label-free quantitation (LFQ), features were extracted from the raw data files using the Minora Feature Detector processing workflow node on Proteome Discoverer. Chromatographic feature-matching was performed with a tolerance of 10 min to align the retention times and enable quantitation of the precursors. No pre-normalization was performed using the Proteome

Discoverer workflow, and raw results were manually normalized using the Median normalization method. A protein was classified as detected if at least two peptides were detected in a minimum of two biological replicates; the mean LFQ across the three replicates was used for further analyses. There was no normalization during protein treatments and peptide injection as described earlier. Normalization between samples was performed only in the data analysis process. Briefly, frequency distributions of Ot protein abundances from six separate runs were plotted (excluded co-purified host proteins). And the median from each plot was used to normalize the whole set, prior to data trimming, making all data set scaled to the same median reference value of 1. Normalized abundance values were then averaged from independent biological replicates (Supplementary table 1). Statistical significance (t-test) was determined using GraphPad Prism software 8.4.3. Subsequently, proteins detected in both IBs and EBs for at least twice were selected for protein abundance comparison, described as Ext/Int ratio. Proteomic data is available from the jPOST repository with the identifier PXD028218.

### Immunoblot analyses

L929 and HUVEC cells were grown in 24-well plates and infected with Ot UT76 at an MOI of 12:1. Non-infectious bacteria were removed by replacing the media 24 hours post-infection. EB and IB populations were prepared four days post-infection. EB was prepared by transferring the cell supernatant to an Eppendorf tube, pelleting at 16,873 g, then resuspending in 100 µl PBS containing 1x protease inhibitor cocktail (ThermoFisher 78425). For TSA56 immunoblots, samples were prepared in the presence of 1 mM iodoacetamide (Sigma-Aldrich catalog number I6125). IB was prepared by removing adherent cells using trypsinization, spinning the cell pellet, and resuspending it in PBS supplemented with protease inhibitors; host cells were lysed using a bead mill for 1 minute. Host cell debris was removed by centrifugation at 500 g for 5 min; the supernatant was transferred to a clean tube, spun at 16,873 g for 5 minutes, and resuspended in 100 µl PBS supplemented with protease inhibitors. Bacterial copy number was quantified by qPCR using *tsa47* primers, as described above. IB and EB samples were diluted to ensure equal loading of bacterial cells on gels. Samples were diluted in 5X SDS loading buffer with and without ß-mercaptoethanol and heated at 95°C for 5 min. Samples were subjected to electrophoresis on a 4-20% gradient gel (Bio-Rad 4561094) and then transferred to a nitrocellulose membrane. Following blocking in PBST containing 5% BSA and 5% skim milk for 30 min, we incubated the blots with primary antibody (1:1,000) in PBST-BSA overnight at 4°C. Membranes were washed and incubated with secondary antibody (1:10,000) conjugated to IRDYE 680RD or IRDYE 800CW (LI-COR Biosciences) in PBST-BSA for 1 hour at room temperature. Membranes were washed, and immunoreactive proteins were visualized using a LiCor Odyssey CLX imager.

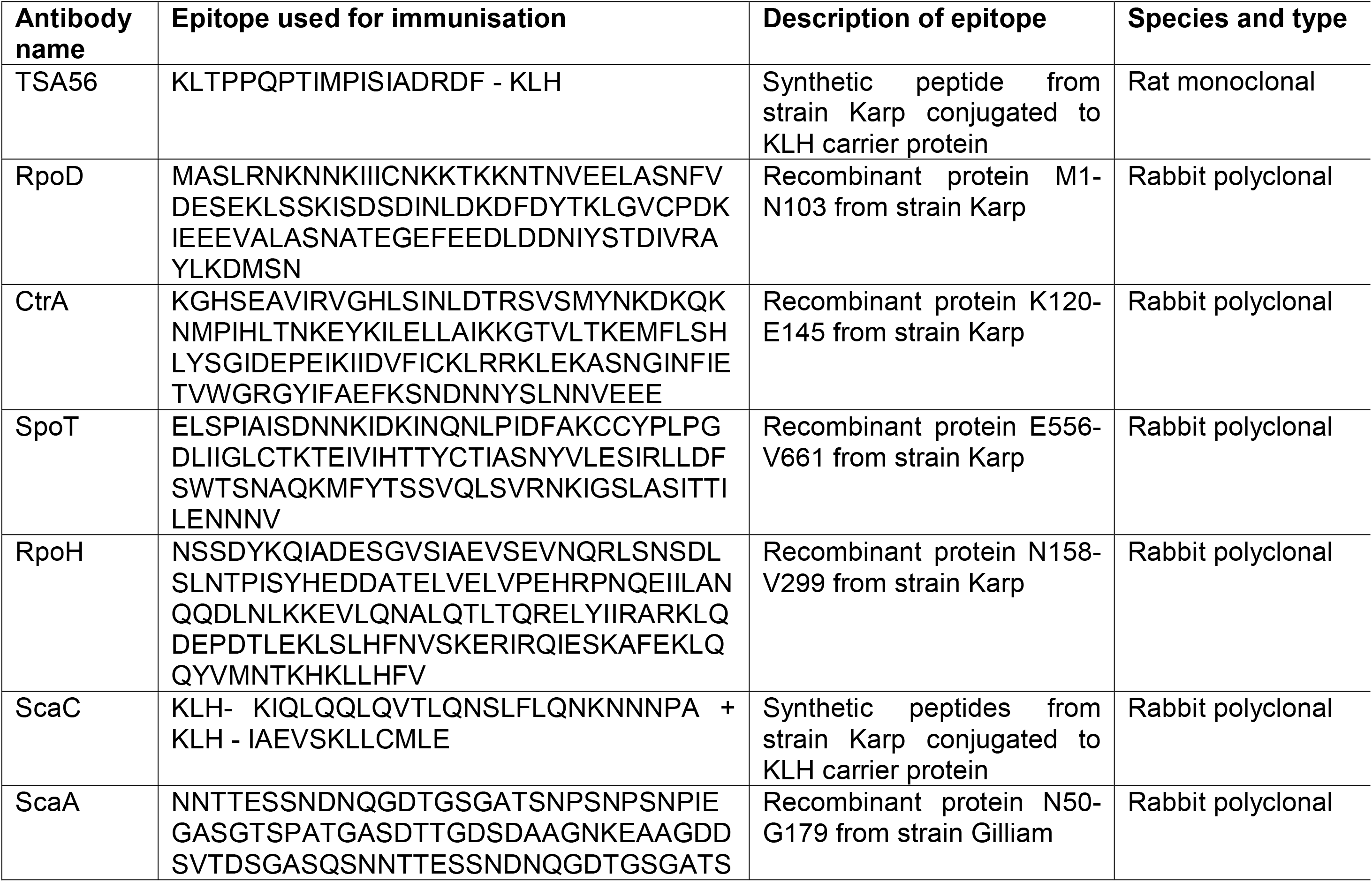
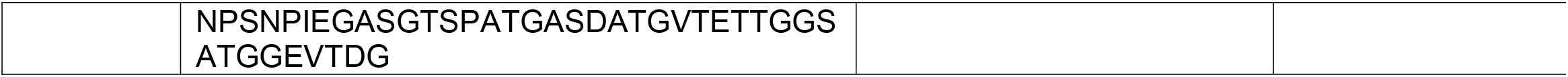

### Quantification of genome copy number per cell

The genome copy number per cell was calculated by determining the number of bacterial genome copies in a sample of purified EB and IB Ot, using *tsa47* qPCR, and dividing by the number of bacterial cells in the same sample, as measured by confocal microscopy. Three µl of purified bacteria were spread evenly across a well of a multi-well slide, dried, and labelled for immunofluorescence microscopy using TSA56, as described above. The bacteria in a selected field were manually counted to determine the number of bacterial cells in the sample.

### Quantification of relative peptidoglycan levels using NOD1 reporter assay

HEK-Blue hNOD1 cells (Invitrogen, hkb-hnod1, USA) were grown in DMEM containing 4.5 g/l glucose, 10% heat-inactivated FBS, 100 U/ml penicillin, 100 mg/ml streptomycin, 100 mg/ml Normocin, 2 mM L-glutamine, and the selective antibiotics blasticidin (30 µg/ml) and Zeocin (100 µg/ml). The cells were seeded on clear-bottom black 96-well plates (Corning 29444-008) 2 days before treatment. Bacterial aliquots from stocks kept in sucrose phosphate glutamine (SPG) at -80°C were heat-inactivated at 90°C for 30 minutes before supplementation to host cells. These studies were carried out in triplicate. After two days, growth media were replaced with HEK-blue detection media (Invivogen hb-det2) for SEAP assessment. The plates were further incubated for six hours at 37°C in a 5% CO_2_ atmosphere and then analyzed spectrophotometrically (Synergy H1, Biotek) at 640 nm. Results were analyzed by GraphPad Prism (GraphPad Software, San Diego California USA), and a t-test was performed to compare bacterial SEAP levels to positive control iE-DAP (γ-D-Glu-mDAP) at 10 µg/ml.

### Growth curve measurements and bead mill treatment

To determine the growth rate of IB and EB Ot from frozen stock were first grown in two T25 flasks containing a monolayer of L929 cells. At 6 days post infection, IB and EB were prepared and used to inoculate fresh L929 cells and the subsequent bacterial growth was measured.

#### EB isolation

the supernatant portion of the culture consisting of EB was transferred to a 50-mL centrifuge tube (Corning 430828) and subjected to a first centrifugation to remove any host cells at 1,500 g for 5 minutes. The supernatant was then transferred into a new 50-ml tube and centrifuged at 3,220 g for 20 minutes to obtain an EB pellet which was resuspended in the fresh media and ready for the next step.

#### IB isolation

fresh media was added directly to the culture flasks containing infected cells and these were disassociated from the adherent surface by using a cell scraper (Corning Falcon, catalog no. 353089). The infected cell suspension was then pipetted into a sterile 2-mL microcentrifuge tube. (Eppendorf 022363352) pre-filled with sterile 0.1-0.15 mm diameter glass beads (Biospec 11079101) before introduced to VWR® Bead Mill Homogenizer (VWR 10158-558) at speed 5 for 1 minutes to break the host cells. Subsequently, the purified IB in the supernatant was separated from host cells debris and glass beads by transferring the supernatant into a new 2-mL microcentrifuge tube and spinning at 300 g for 5 minutes. The bacterial concentration in both EB and IB suspension was quantified by qPCR as described above, and EB and IB Ot was infected onto a fresh monolayer of L929 cells in 12-well plates (Thermo Scientific 150628). Bacteria were added at 2.83E+04 bacterial copies/well which corresponds to an MOI of ∼1:10. The number of intracellular bacteria was then measured at 3, 24, 96 and 168 hours post infection by *tsa47* qPCR.

#### Physical stress induced by bead mill

IB and EB isolation and infection were performed as described above and then exposed to physical stress with 1 or 2 minutes agitation in the bead mill. Bacteria were then infected onto L929 cells at a concentration of 1.00E+06 copies/well (MOI = 2:1). The quantity of intracellular bacteria was then determined at 3, 24, 96, 120, 144 and 168 hours post infection by qPCR.

#### Osmotic stress induced by water treatment

Quantification of the effect of osmotic shock on Ot viability was performed in the same way as for physical stress except purified bacteria were resuspended in pure water for 5 minutes prior to infecting host cells. Bacterial copy number after 7 days was quantified by qPCR.

##### Entry inhibition experiments

IB and EB forms of Ot were isolated from infected host cells as described above. To assess the impact of different interventions on bacterial entry, IB and EB form Ot were added to uninfected L929 or HUVEC cells, labelled with HPG and fixed three hours post infection. The number of bacteria inside host cells was counted manually with at least 100 host cells counted for each condition/replicate. To measure the role of clathrin, 0.2 µM siRNA (clathrin, catalog number 4390825 or scramble catalog number 4390843, ThermoFisher Scientific) was added 1 day prior to infection. To measure the effect of perturbing clathrin-mediated endocytosis or macropinocytosis, inhibitors were added for one hour prior to infection and kept in the media during the three hours of infection. Dynasore hydrate (inhibits endocytosis, catalog number D7693, Sigma) was used at 100 µM, amiloride (inhibit micropinocytosis, 5-(N-Ethyl-N-isopropyl, catalog number A3085, Sigma) at 50 µM and zoniporide hydrochloride hydrate (inhibit micropinocytosis, catalog number SML0076, Sigma) at 100 µM. In order to determine the role of bacterial surface proteins TSA56 and ScaC, IB and EB Ot were added to uninfected host cells at the same time as antibodies against TSA56 and ScaC and the number of intracellular bacteria at three hours post infection quantified by microscopy.

### Statistical analyses

All statistical analyses were carried out using GraphPad Prism version 9.2.0. Comparisons between two groups with >10 datapoints per group were carried out using a students unpaired t test, whilst comparisons between two groups with ≤10 datapoints per group were carried out using a Mann-Whitney test. Comparisons between more than two groups were carried out using one-way ANOVA followed by Dunnetts comparisons. Ns = no significant difference detected, P > 0.05; * P ≤ 0.05; ** P ≤ 0.01; *** P ≤ 0.001; **** P ≤ 0.0001.

## Acknowledgements and Funding

We thank each laboratory member for valuable discussions and support. We are grateful to the following people for providing helpful feedback and insightful comments related to the project and the manuscript: Jonathan Dworkin from Columbia University, Dave Dubnau from PHRI, and Diana Stafforini from Life Science Editors. We also appreciate the support from Patricia Greenberg and Tracy Andrews at the Rutgers University Biostatistics and Epidemiology Services (RUBIES).

This work was funded by a Royal Society Dorothy Hodgkin Research Fellowship (JS) (DH140154), a Newton Fund MRC-NSTDA grant (YJ/JS) (MR/N012380/1), and National Institute of Health/National Institute of Allergy and Infectious Diseases grants (R21AI144385 and R56AI148645) (JS), and A*STAR core funding and Singapore National Research Foundation under its NRF-SIS “SingMass” scheme (RMS).

## Data availability

Source data are provided with this paper as raw data files. Proteomics data is available on ProteomeXchange (PXD028218) and jPOST (JSPT001299)^28^.

## Notes

### Competing Interest Statement

The authors have declared no competing interest.

### Summary of Updates

New data in Fig. 2C, 3H, Supp Fig 9

## References

1. Taylor, A., Paris, D. & Newton, P. A Systematic Review of Mortality from Untreated Scrub Typhus (Orientia tsutsugamushi). PLoS Negl Trop Dis 9, e0003971, doi:10.1371/journal.pntd.0003971 (2015).

2. Luce-Fedrow, A. et al. A Review of Scrub Typhus (Orientia tsutsugamushi and Related Organisms): Then, Now, and Tomorrow. Tropical Medicine and Infectious Disease 3, 8, doi:10.3390/tropicalmed3010008 (2018).

3. McClure, E. et al. Engineering of obligate intracellular bacteria: progress, challenges and paradigms. Nat Rev Microbiol, doi:10.1038/nrmicro.2017.59 (2017).

4. Salje, J. Cells within cells: Rickettsiales and the obligate intracellular bacterial lifestyle. Nat Rev Microbiol, doi:10.1038/s41579-020-00507-2 (2021).

5. Rikihisa, Y. & Ito, S. Localization of electron-dense tracers during entry of Rickettsia tsutsugamushi into polymorphonuclear leukocytes. Infect Immun 30, 231–243 (1980).

6. Lee, J. et al. Fibronectin facilitates the invasion of Orientia tsutsugamushi into host cells through interaction with a 56-kDa type-specific antigen. J Infect Dis 198, 250–257, doi:10.1086/589284 (2008).

7. Ha, N., Cho, N., Kim, Y., Choi, M. & Kim, I. An autotransporter protein from Orientia tsutsugamushi mediates adherence to nonphagocytic host cells. Infect Immun 79, 1718–1727, doi:10.1128/IAI.01239-10 (2011).

8. Ha, N. et al. Immunization with an Autotransporter Protein of Orientia tsutsugamushi Provides Protective Immunity against Scrub Typhus. PLoS Negl Trop Dis 9, e0003585, doi:10.1371/journal.pntd.0003585 (2015).

9. Chu, H. et al. Exploitation of the endocytic pathway by Orientia tsutsugamushi in nonprofessional phagocytes. Infect Immun 74, 4246–4253, doi:10.1128/IAI.01620-05 (2006).

10. Kim, H., Choi, M. & Kim, I. Role of Syndecan-4 in the cellular invasion of Orientia tsutsugamushi. Microb Pathog 36, 219–225, doi:10.1016/j.micpath.2003.12.005 (2004).

11. Cho, B., Cho, N., Seong, S., Choi, M. & Kim, I. Intracellular invasion by Orientia tsutsugamushi is mediated by integrin signaling and actin cytoskeleton rearrangements. Infect Immun 78, 1915–1923, doi:10.1128/IAI.01316-09 (2010).

12. Giengkam, S., et al. Improved Quantification, Propagation, Purification and Storage of the Obligate Intracellular Human Pathogen Orientia tsutsugamushi. PLoS Negl Trop Dis 9, e0004009, doi:10.1371/journal.pntd.0004009 (2015).

13. Atwal, S. et al. Evidence for a peptidoglycan-like structure in Orientia tsutsugamushi. Mol Microbiol 105, 440–452, doi:10.1111/mmi.13709 (2017).

14. Min, C. et al. Genome-based construction of the metabolic pathways of Orientia tsutsugamushi and comparative analysis within the Rickettsiales order. Comp Funct Genomics 2008, 623145, doi:10.1155/2008/623145 (2008).

15. Atwal, S. et al. Clickable methionine as a universal probe for labelling intracellular bacteria. J Microbiol Methods 169, 105812, doi:10.1016/j.mimet.2019.105812 (2020).

16. Rikihisa, Y. Anaplasma phagocytophilum and Ehrlichia chaffeensis: subversive manipulators of host cells. Nat Rev Microbiol 8, 328–339, doi:10.1038/nrmicro2318 (2010).

17. Elwell, C., Mirrashidi, K. & Engel, J. Chlamydia cell biology and pathogenesis. Nat Rev Microbiol 14, 385–400, doi:10.1038/nrmicro.2016.30 (2016).

18. Otten, C., Brilli, M., Vollmer, W., Viollier, P. & Salje, J. Peptidoglycan in obligate intracellular bacteria. Mol Microbiol 107, 142–163, doi:10.1111/mmi.13880 (2018).

19. Jacquier, N., Frandi, A., Pillonel, T., Viollier, P. & Greub, G. Cell wall precursors are required to organize the chlamydial division septum. Nat Commun 5, 3578, doi:10.1038/ncomms4578 (2014).

20. Irving, S. E., Choudhury, N. R. & Corrigan, R. M. The stringent response and physiological roles of (pp)pGpp in bacteria. Nat Rev Microbiol, doi:10.1038/s41579-020-00470-y (2020).

21. Batty EM, C. S., Blacksell SB, Richards, A, Paris D, Bowden, R, Chan C, Lachumanan R, Day N, Donnelly P, Chen SL, Salje J. Long-read whole genome sequencing and comparative analysis of six strains of the human pathogen Orientia tsutsugamushi. Plos Negl Trop Dis (2018).

22. Mika-Gospodorz, B. et al. Dual RNA-seq of Orientia tsutsugamushi informs on host-pathogen interactions for this neglected intracellular human pathogen. Nat Commun 11, 3363, doi:10.1038/s41467-020-17094-8 (2020).

23. Joshi, K. K. & Chien, P. Regulated Proteolysis in Bacteria: Caulobacter. Annu Rev Genet 50, 423–445, doi:10.1146/annurev-genet-120215-035235 (2016).

24. Lamason, R. et al. Rickettsia Sca4 Reduces Vinculin-Mediated Intercellular Tension to Promote Spread. Cell 167, 670–683.e610, doi:10.1016/j.cell.2016.09.023 (2016).

25. Lee, S. Optimal integration of wide field illumination and holographic optical tweezers for multimodal microscopy with ultimate flexibility and versatility. Opt Express 26, 8049–8058, doi:10.1364/OE.26.008049 (2018).

26. Huang, B., Wang, W., Bates, M. & Zhuang, X. Three-dimensional super-resolution imaging by stochastic optical reconstruction microscopy. Science 319, 810–813, doi:10.1126/science.1153529 (2008).

27. Wang, Y. et al. Localization events-based sample drift correction for localization microscopy with redundant cross-correlation algorithm. Opt Express 22, 15982–15991, doi:10.1364/OE.22.015982 (2014).

28. Okuda, S. et al. jPOSTrepo: an international standard data repository for proteomes. Nucleic Acids Res 45, D1107–D1111, doi:10.1093/nar/gkw1080 (2017).

