## Supplementary Figures for "The obligate intracellular bacterium *Orientia tsutsugamushi* differentiates into a developmentally distinct extracellular state"

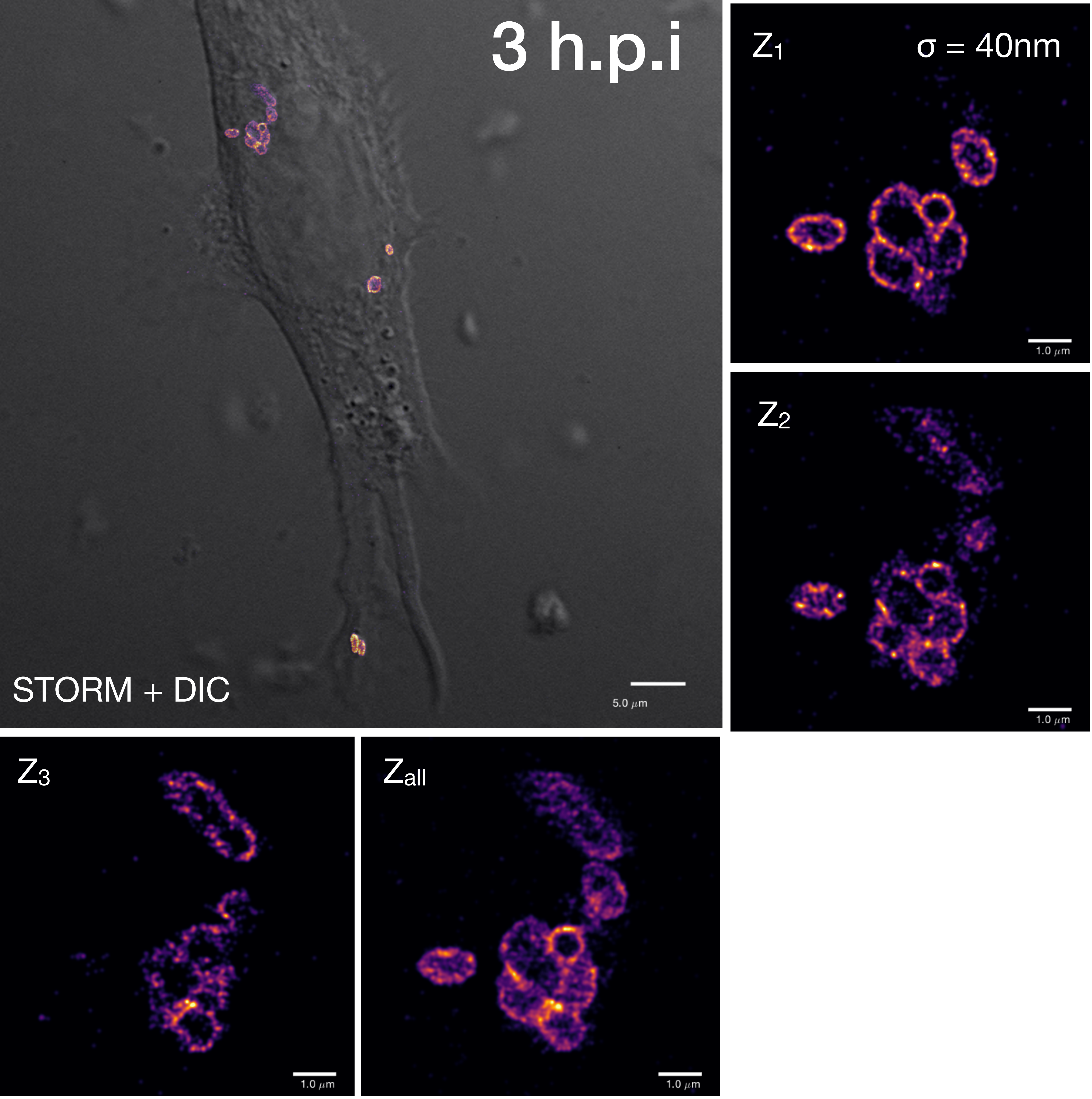


**Supplementary Figure 1.** Representative STORM images of intracellular and extracellular Ot at different times after infection. Extracellular bacteria are shown with an arrow except on 7 d.p.i. where a full field of extracellular bacteria is shown. H.p.i. = hours post infection, d.p.i. = days post infection. Ot strain UT76 in L929 cells.

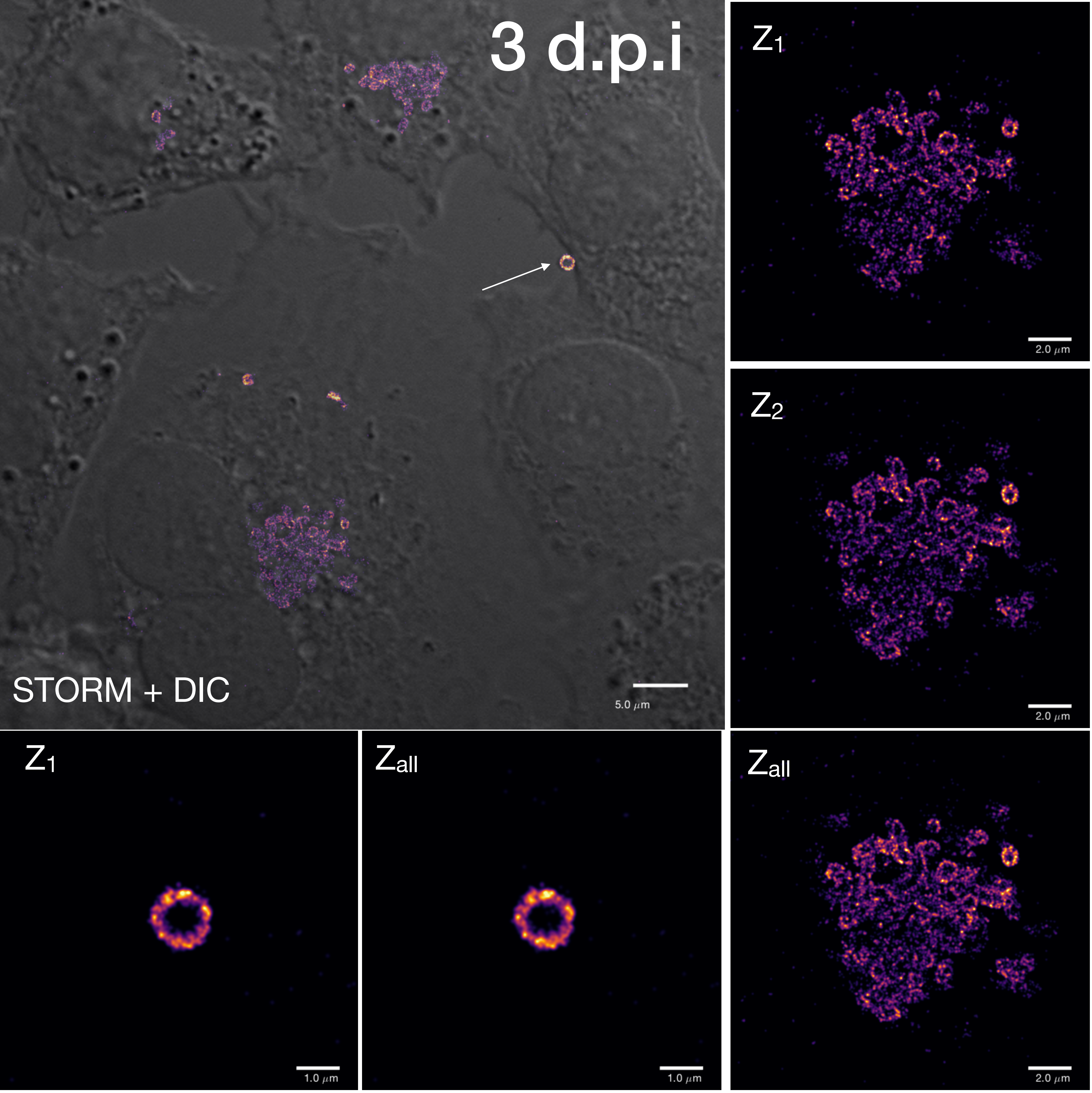

3 d.p.i
STORM + DIC
5.0 μm
Z1
2.0 μm
Z2
2.0 μm
Z1
1.0 μm
Zall
1.0 μm
Zall
2.0 μm

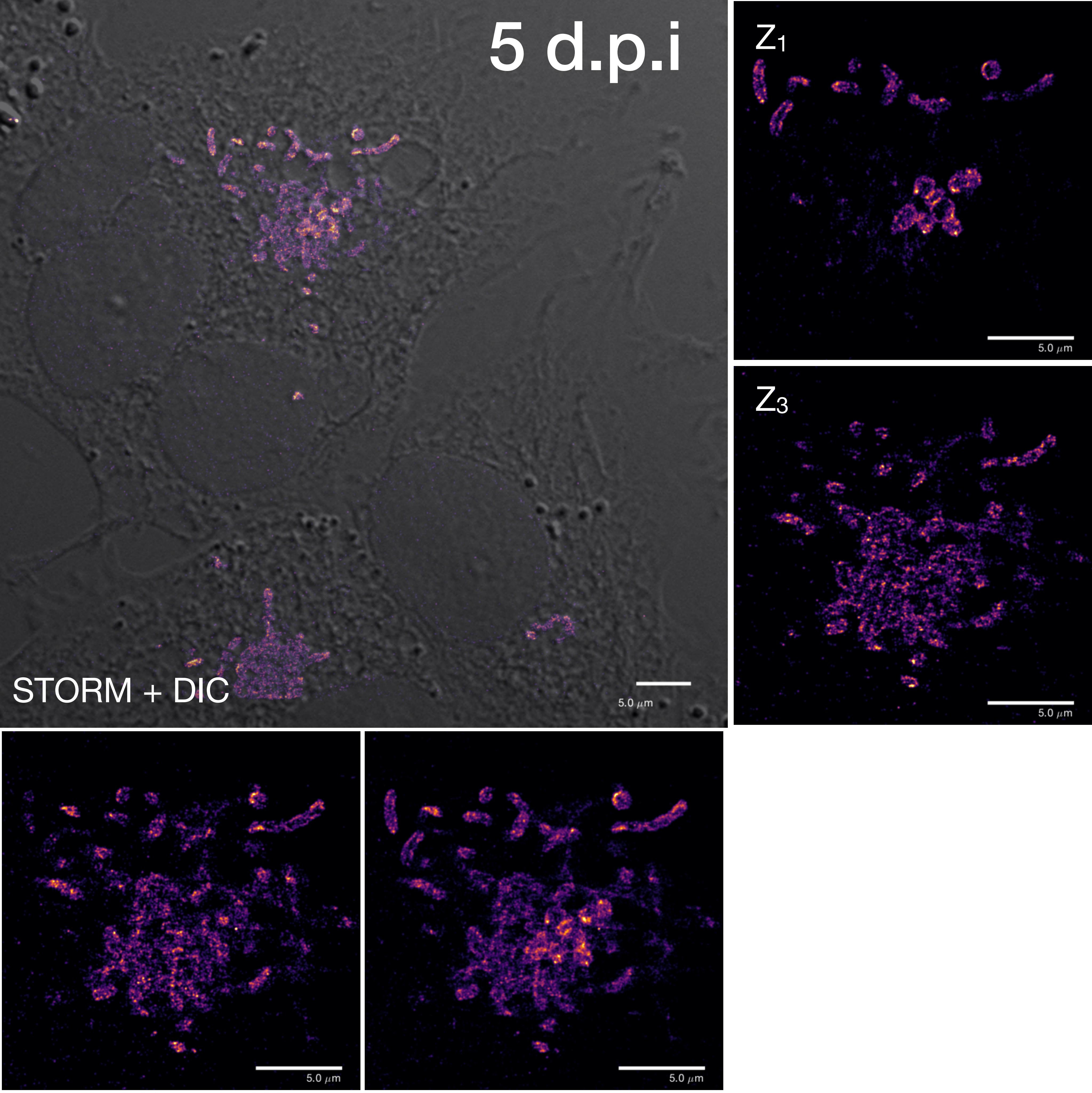

5 d.p.i
STORM + DIC
5.0 μm
Z1
5.0 μm
Z3
5.0 μm
5.0 μm
5.0 μm

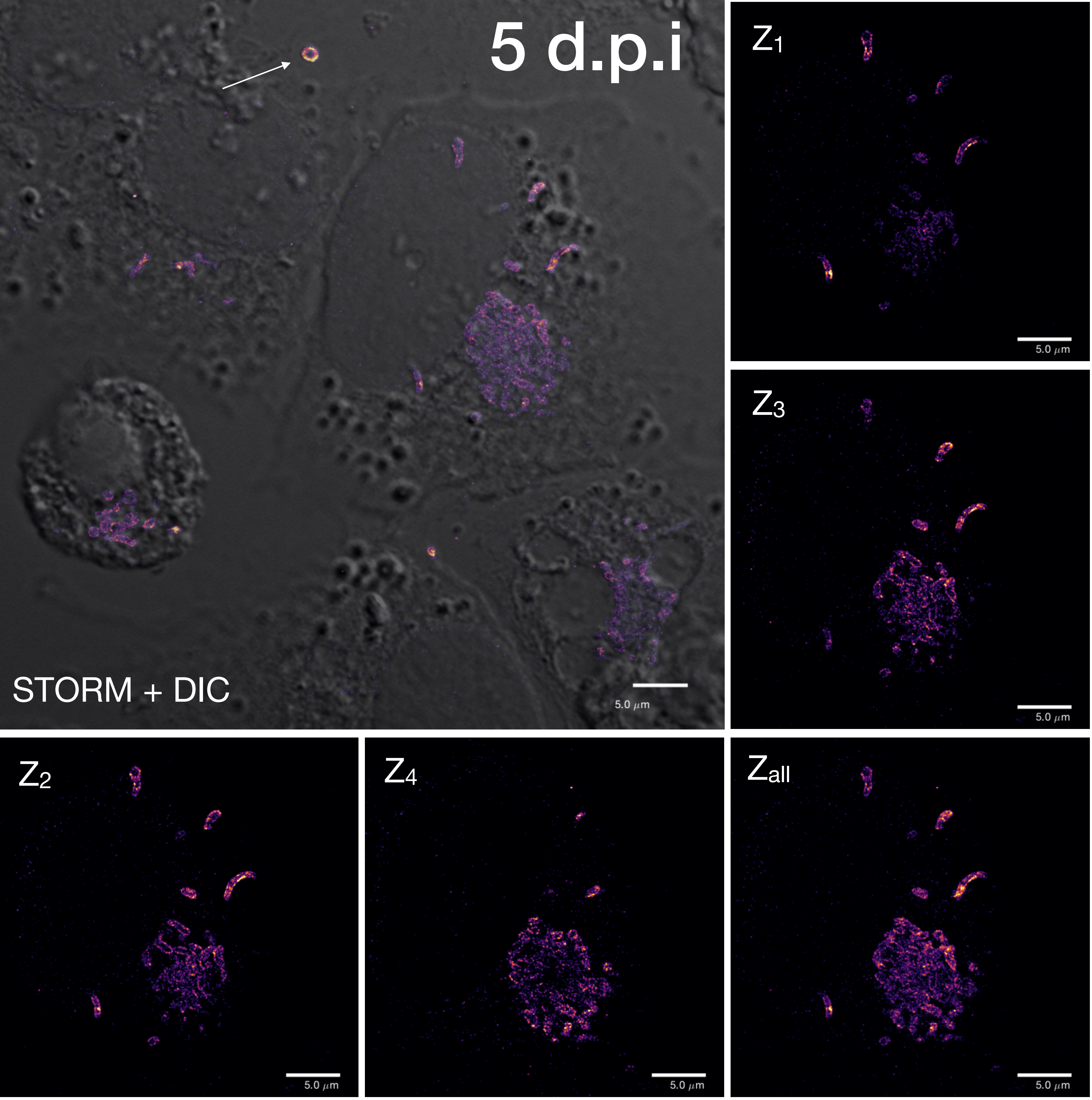
5 d.p.i
STORM + DIC
5.0 µm
Z1
5.0 µm
Z3
5.0 µm
Z2
5.0 µm
Z4
5.0 µm
Zall
5.0 µm

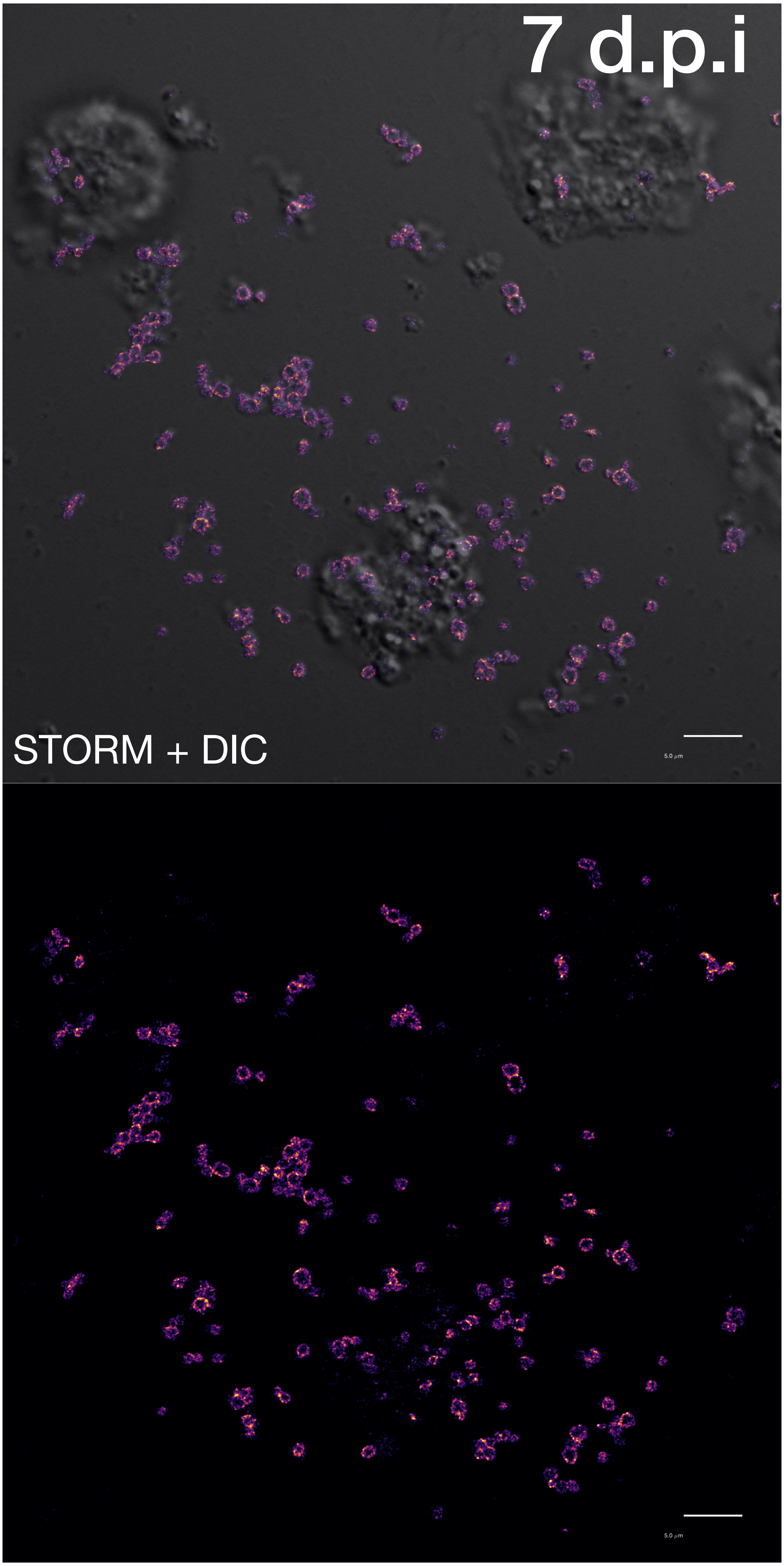
7 d.p.i
STORM + DIC
5.0 μm
5.0 μm

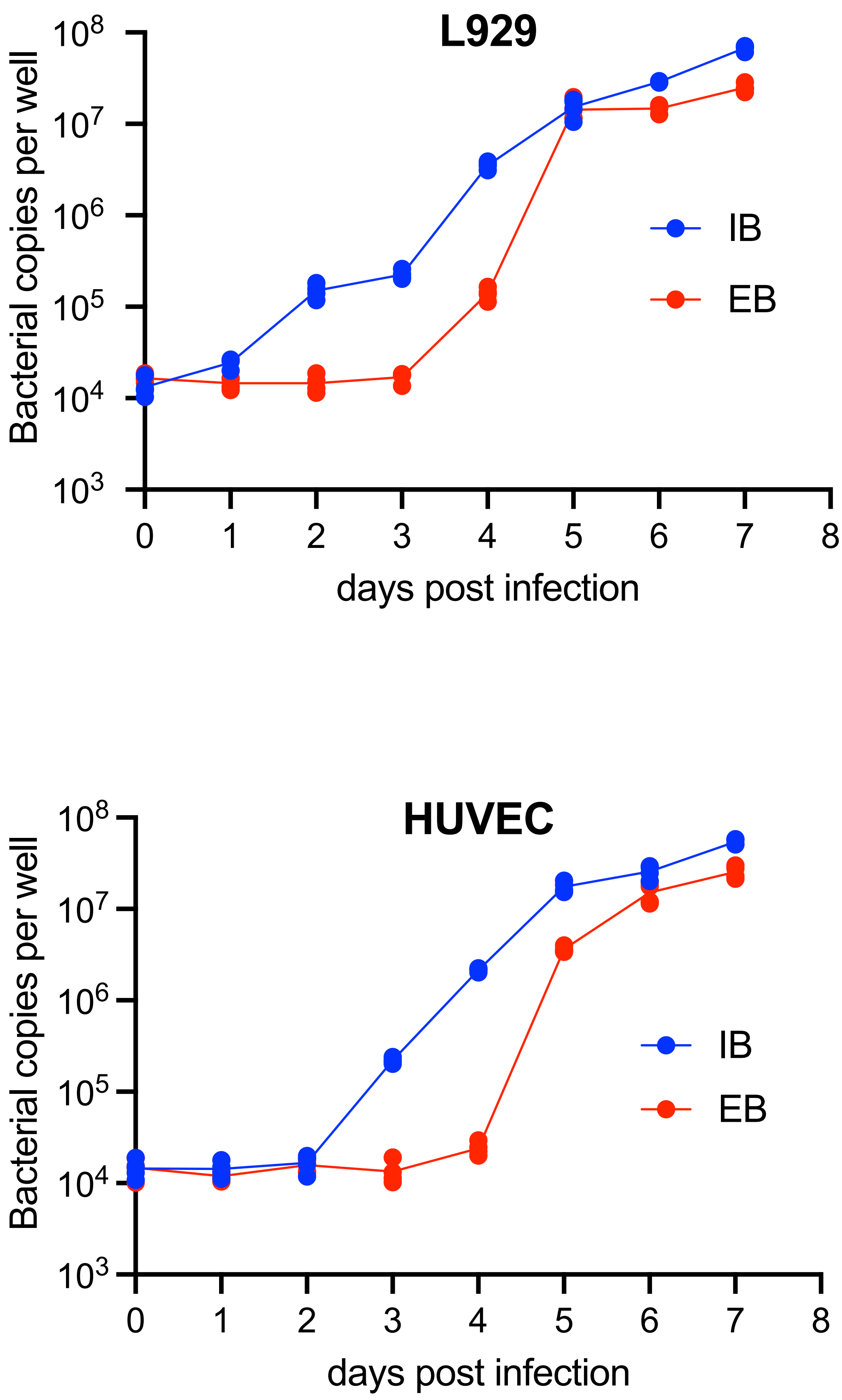


**Supplementary Figure 2. Ot is released into the supernatant at high levels after five days of infection.** Growth curve measuring the number of Ot bacteria in the intracellular (IB) and extracellular (EB) fraction. L929 and HUVEC cells were grown in 24 well plates and infected with frozen Ot. Each day four replicate wells were analyzed. The entire supernatant was collected to generate the EB fraction, and the remaining adhered cells were analyzed as the IB fraction. Ot DNA was extracted and the levels of bacteria measured using qPCR with primers against the single copy gene *tsa47.*

A.

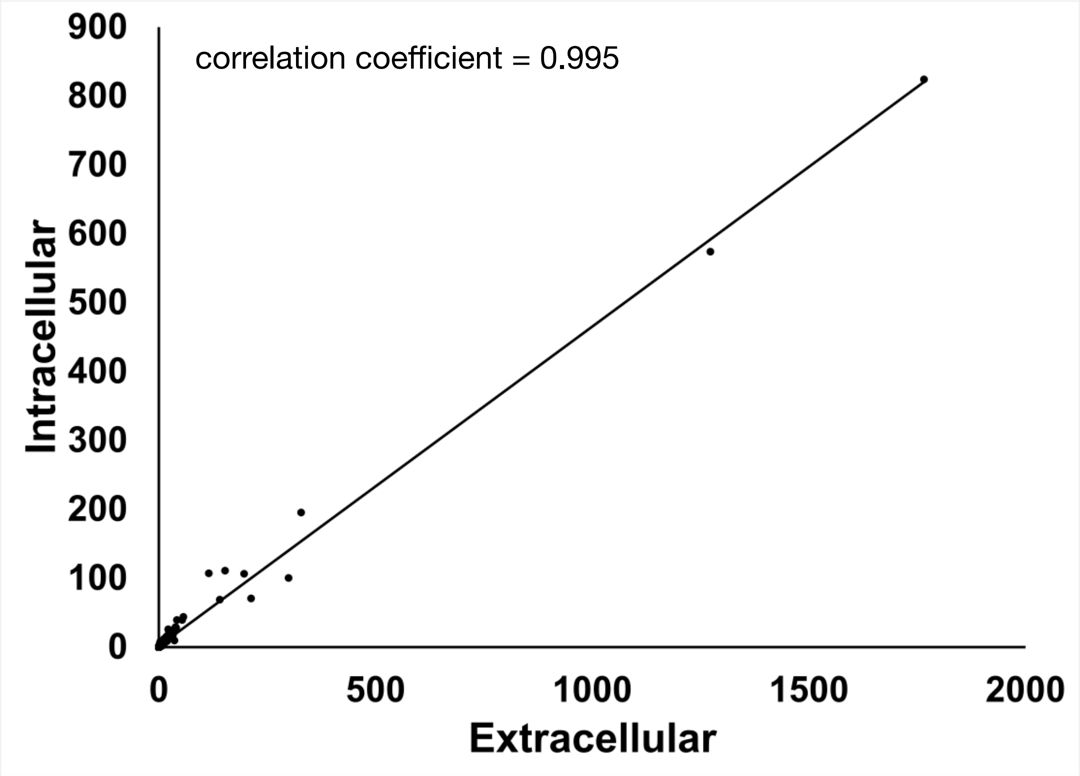


B.

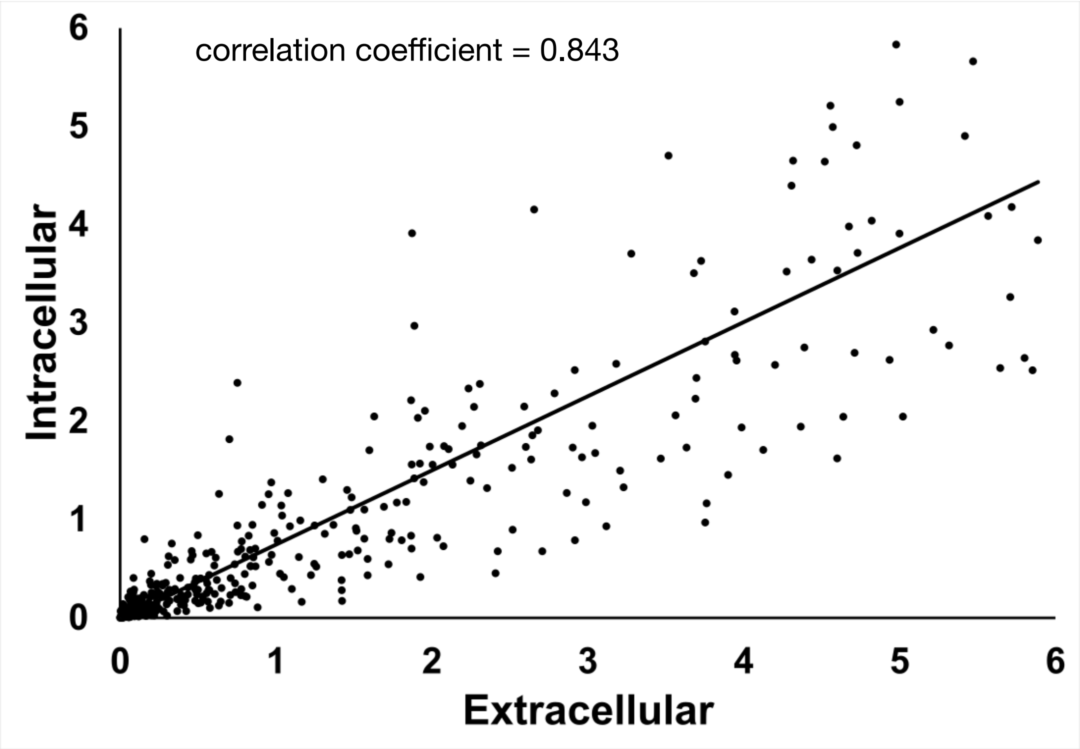


**Supplementary Figure 3. A comparison of the protein profiles of IB and EB forms of Ot.** Bacteria were grown in L929 cells at an MOI of 410:1 and harvested four days post infection. IB and EB Ot were isolated and analyzed by shotgun proteomics. A comparison of relative levels of proteins using normalized LFQ in IB and EB is shown here. B. shows a subset of the data in A, whereby highly abundant proteins have been removed from the analysis.

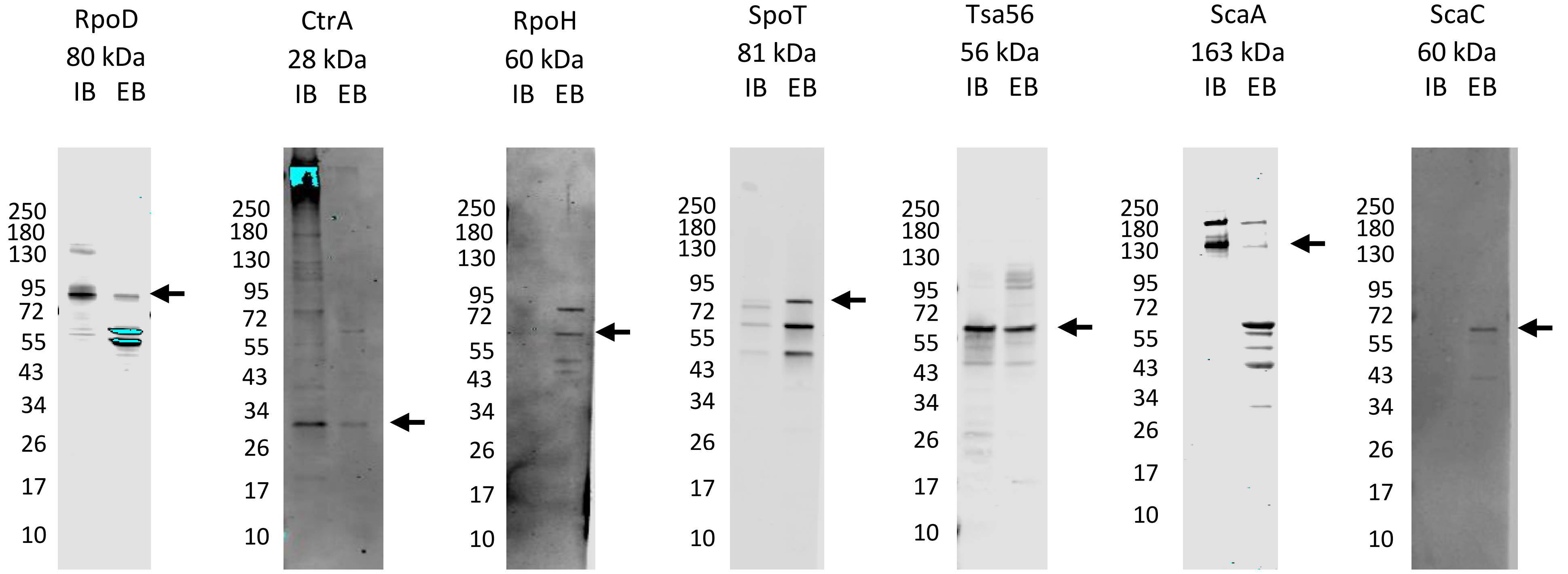


**Supplementary Figure 4. Immunoblot showing specificity of antibodies used in this study.** Ot cell lysate was boiled in SDS-PAGE loading buffer, separated by SDS-PAGE and analysed by immunoblot using antibodies against RpoD, CtrA, RpoH, SpoT, TSA56, ScaA and ScaC. IB = intracellular Ot, EB = extracellular Ot.

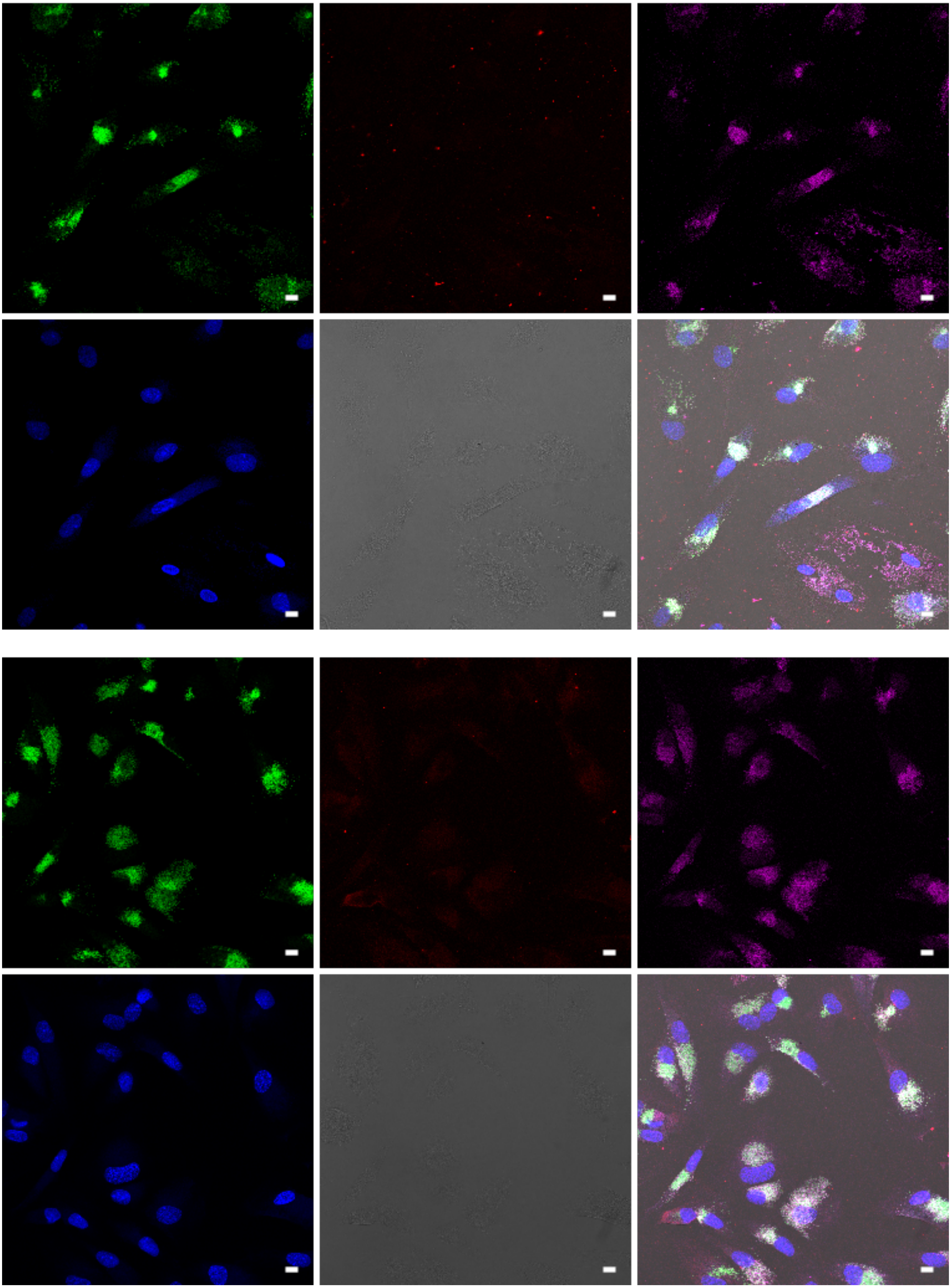

**Supplementary Figure 5. Distinct patterns of expression of ScaA and ScaC are shown by immunofluorescence confocal microscopy.** HUVEC cells were infected with Ot for five days and labelled for DNA (Hoechst, blue), translational activity (HPG, green), ScaA (anti-ScaA antibody, magenta) or ScaC (anti-ScaC antibody, red). Scale bar = 10 µm. Images from two independent experiments are shown here.

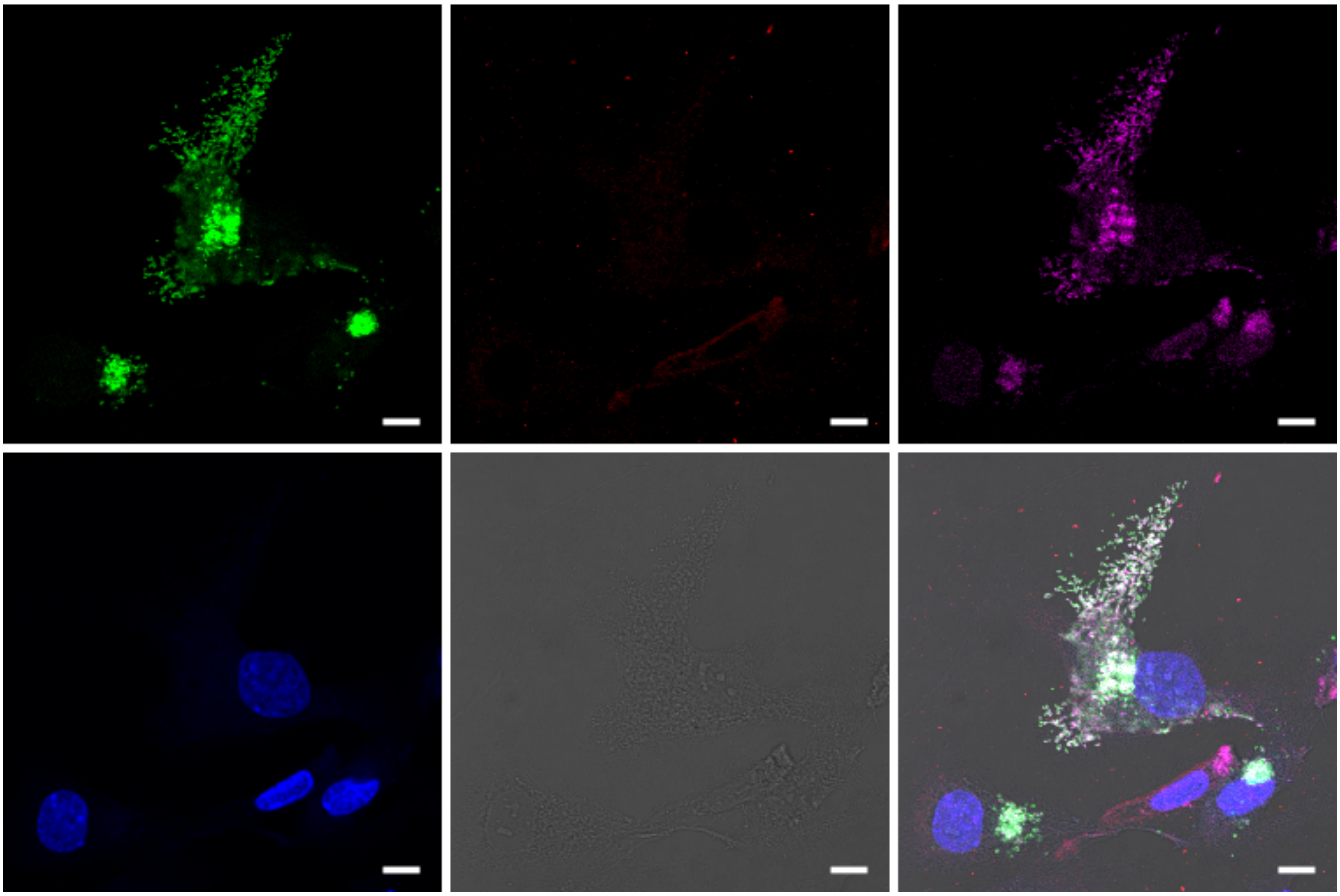

**Supplementary Figure 5. cont.**

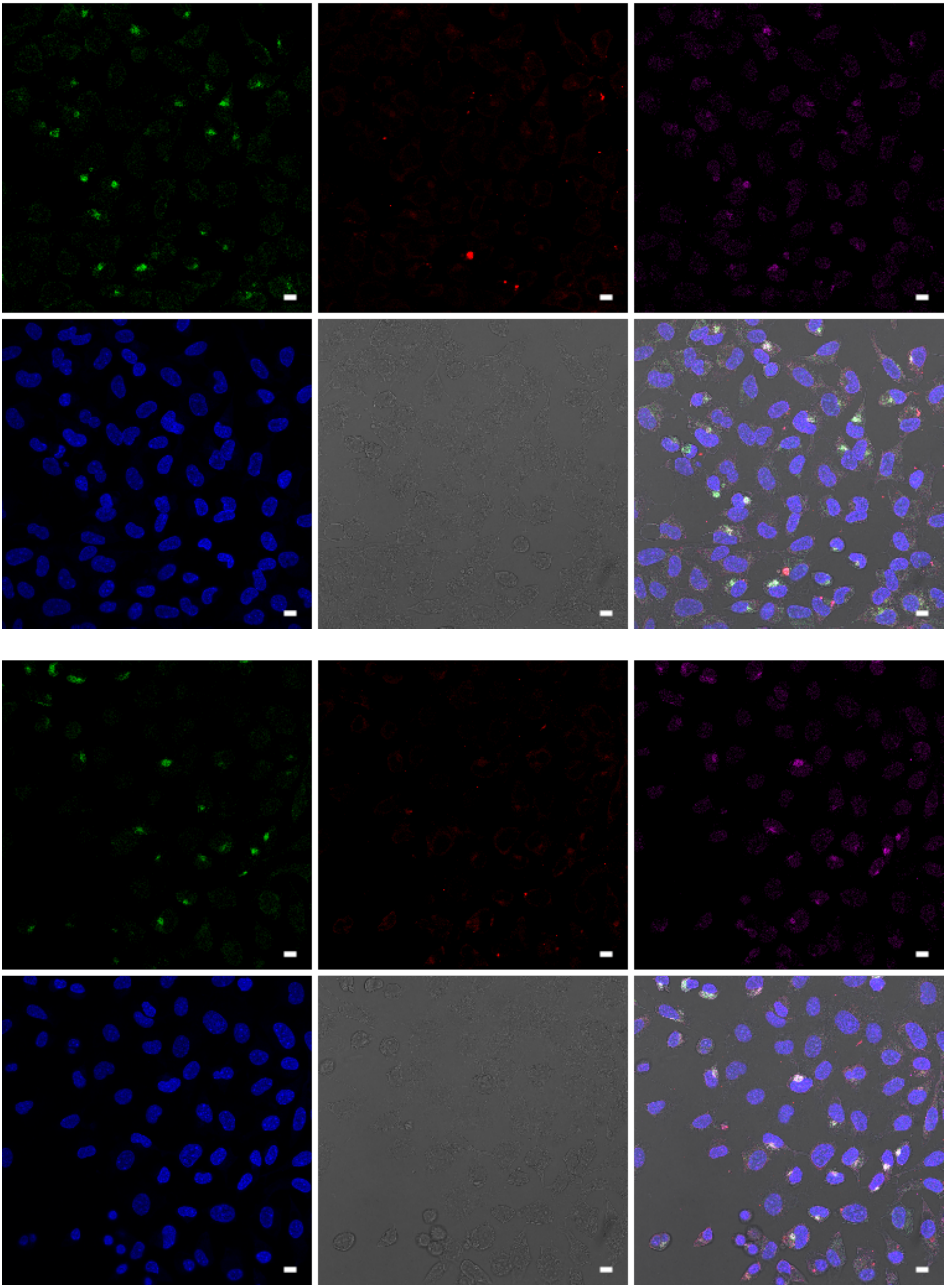

**Supplementary Figure 6. Distinct patterns of expression of ScaA and ScaC are shown by immunofluorescence confocal microscopy.** L929 cells were infected with Ot for four days and labelled for DNA (Hoechst, blue), translational activity (HPG, green), ScaA (anti-ScaA antibody, magenta) or ScaC (anti-ScaC antibody, red). Scale bar = 10 µm. Images from two independent experiments are shown here.

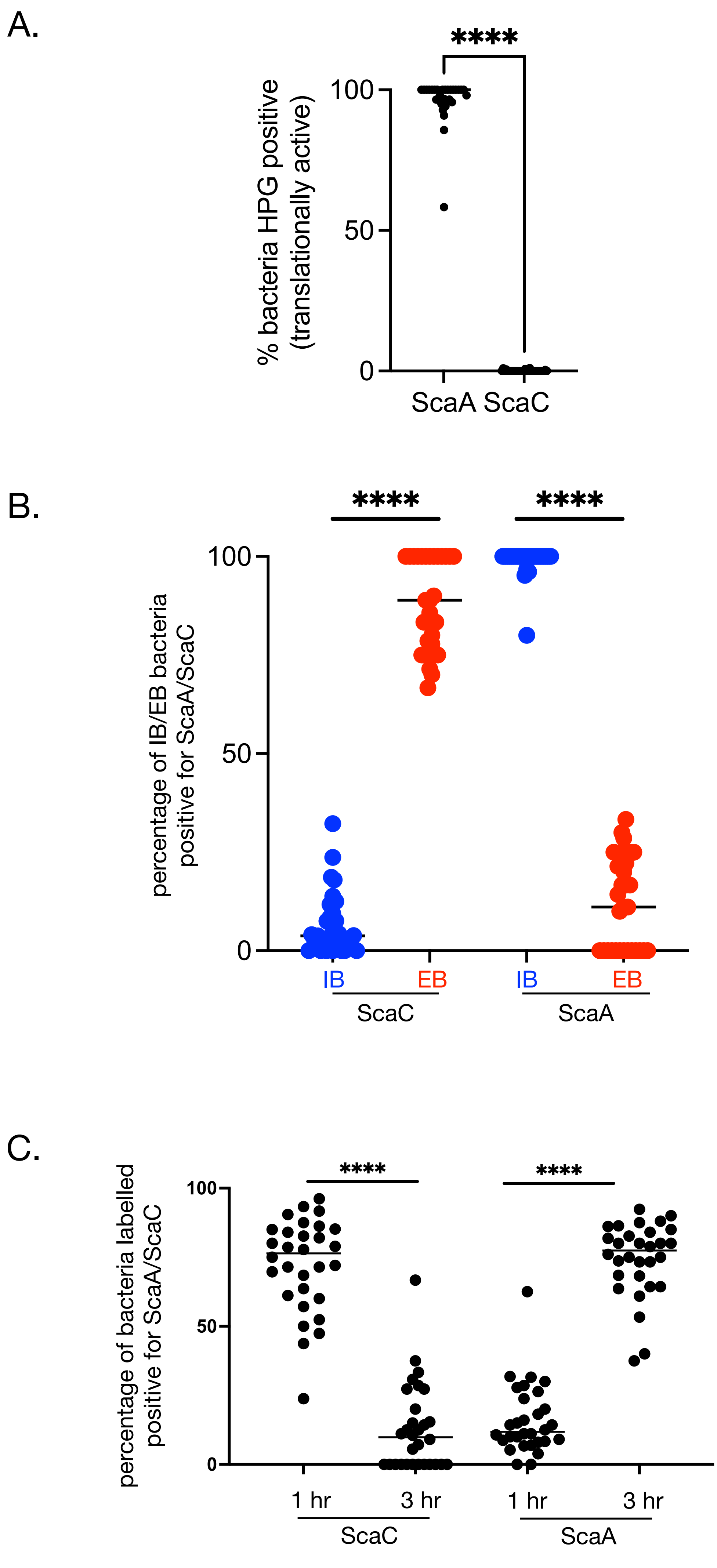


**Supplementary Figure 7: Autotransporter proteins ScaA and ScaC are present at different levels on IB and EB forms of Ot in L929.**
A.Quantification of translational activity, as determined by a positive confocal microscopy signal observed with the clickable methionine probe L-homopropylgylglycine (HPG). L929 cells were infected with Ot for five days, then fixed and labelled with HPG and anti-ScaA or anti-ScaC antibodies. ScaA/ScaC-positive bacteria were scored as being positive or negative for HPG labelling. B. Quantification of confocal micrographs of L929 cells infected with Ot and labelled as in Supp. Fig. 6. The percentage of IB and EB Ot at five days post-infection that were positively labelled with anti-ScaA and -ScaC antibodies was counted. C. Quantification of ScaA- and ScaC-positive IB bacteria inside L929 cells one and three hours post-infection, showing an increase in ScaA labelling and a decrease in ScaC labelling. Throughout this figure quantification was carried out by randomly imaging 30 fields across three independent experiments. Graphs show individual values and means. Statistical significance was calculated using an unpaired t test with Graphpad Prism software. * p-value ≤ 0.05, ** p-value ≤ 0.01, *** p-value ≤ 0.001, **** p-value ≤ 0.0001.

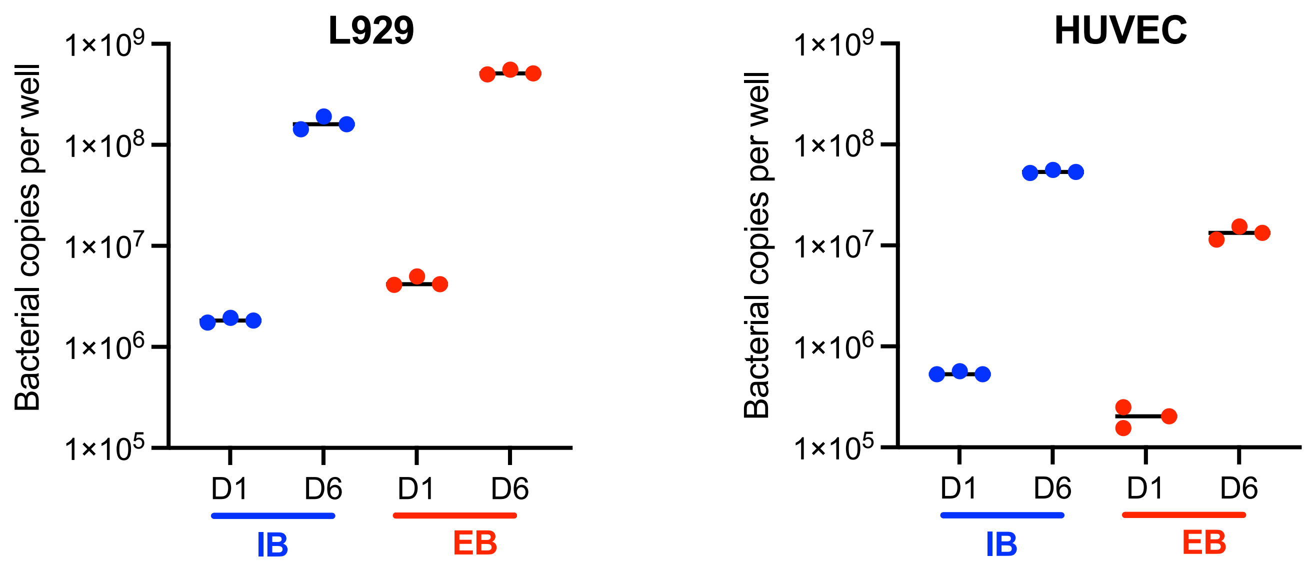


**Supplementary Figure 8. Growth of IB and EB form Ot in L929 and HUVEC cells.** IB and EB Ot were isolated from infected L929 cells five days post-infection and the same inoculum grown in fresh L929 or HUVEC cells in a 24-well plate. The number of intracellular bacteria was then quantified at 1 and 6 days post-infection using qPCR against *tsa47,* a single copy Ot gene*.* Doubling times of IB and EB Ot were similar within each cell type. D1 = day 1 post infection; D6 = day 6 post infection.

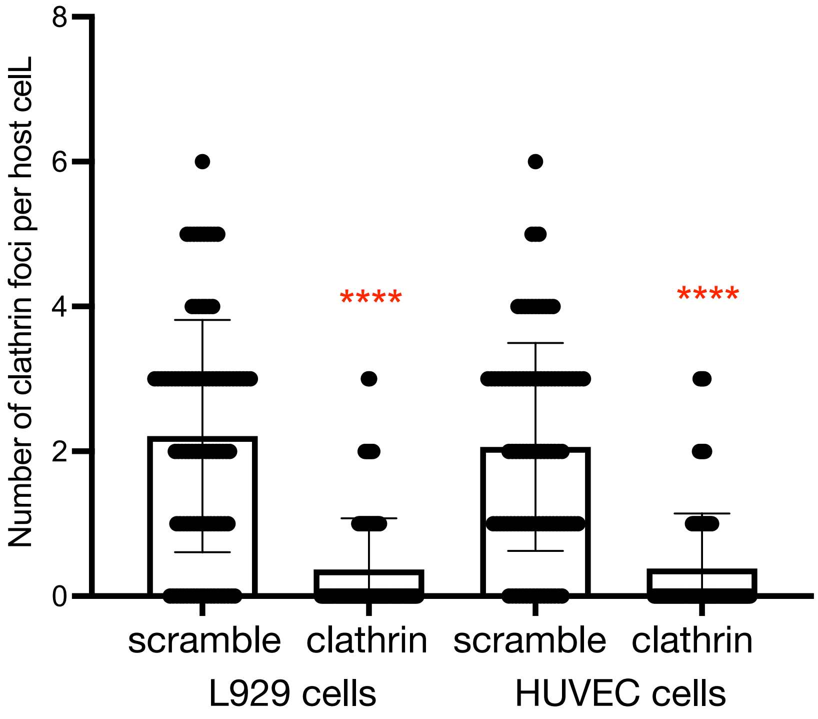


**Supplementary Figure 9. Clathrin siRNA reduces clathrin levels in L929 and HUVEC cells.** Data showing that siRNA targeting clathrin reduces the number of clathrin foci in uninfected L929 and HUVEC cells compared with with a scramble siRNA control. Cells were treated with 0.2 µM siRNA for one hour then fixed and labelled with an anti-clathrin antibody. The number of clathrin foci was enumerated for 100 host cells. Statistical significance was determined using an unpaired t test using GraphPad Prism. .**** p ≤ 0.0001.

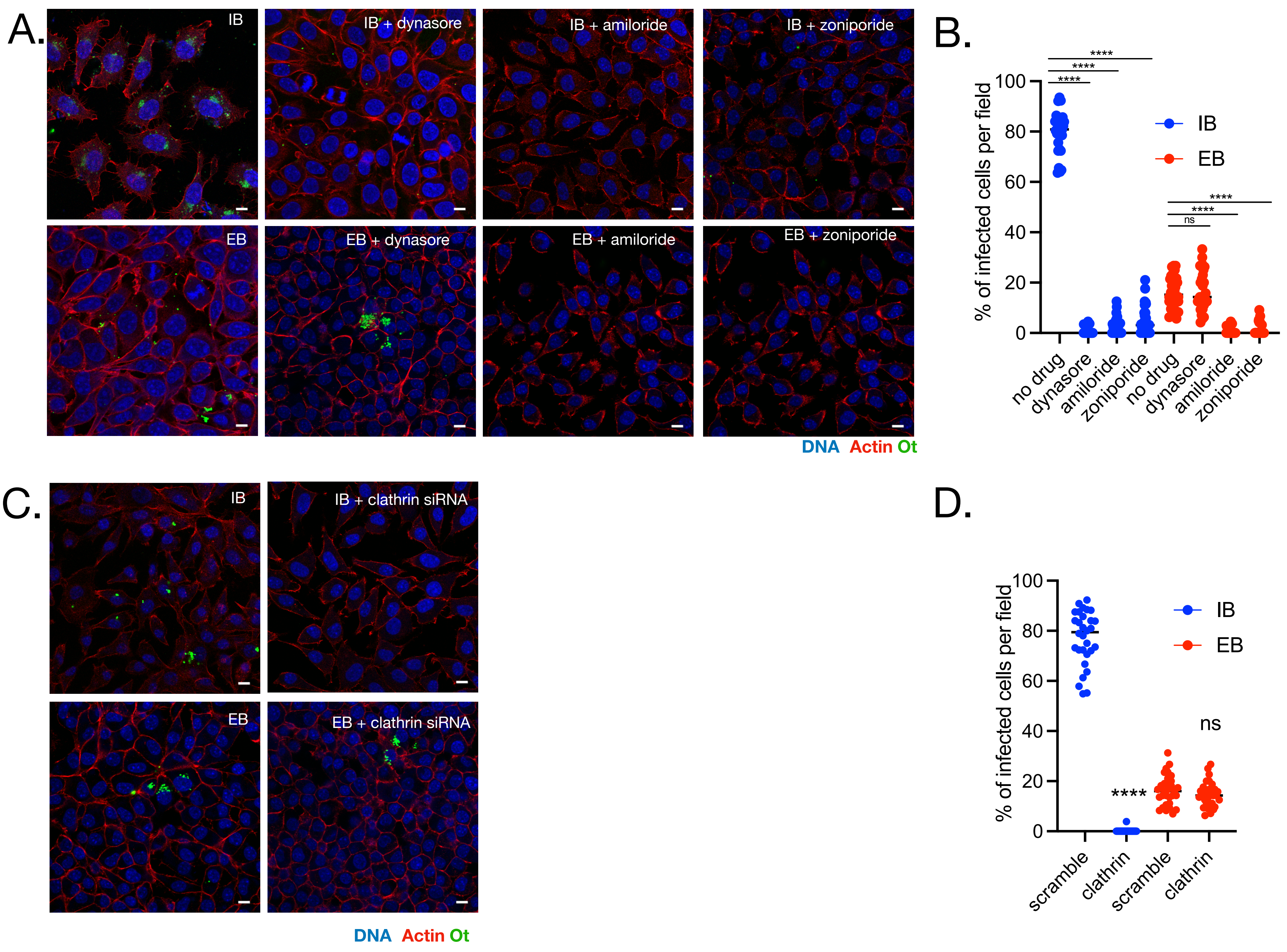


**Supplementary Figure 10. Entry of IB and EB Ot into L929 cells.** A. Confocal microscopy images and (B.) quantification of the effect of clathrin-mediated endocytosis and macropinocytosis inhibitors on entry of IB and EB Ot. IB and EB were isolated from L929 cells 5 days post infection and then seeded onto fresh L929 cells that had been treated with an inhibitor of clathrin-mediated endocytosis (100 µM dynasore) or macropinocytosis (50 µM amiloride or 100 µM zoniporide). Cells were fixed three hours post-infection. Dynasore inhibits entry of IB but not EB, indicating that clathrin-mediated endocytosis is the major mode of entry of this population, whilst amiloride and zoniporide inhibited entry of both IB and EB. This observation points to a role for macropinocytosis, but not clathrin-mediated endocytosis, in EB's entry into mammalian cells. Red = actin (phalloidin), blue = Hoechst (DNA), green = Ot (anti-TSA56 antibody). Scale bar = 10 µm. Images (C) and quantification (D) of the effect of 0.2 µM siRNA against clathrin on entry of EB and IB Ot into L929 cells. Experiments were carried out as in A. and B. Quantification of B. and D. were carried out as follows: 30 fields were randomly imaged over 3 independent experiments (10 fields per experiment). In each field the percentage of infected host cells was counted. Individual values and means are shown here. Statistical significance in (B) was determined by comparing all groups using a one-way ANOVA, followed by a Dunnett's multiple comparison tests with the control, and in (D.) by performing a Mann Whitney test comparing the scramble control and the clathrin siRNA treated condition for IB and EB separately, using GraphPad Prism software. **** $p \leq 0.0001$; ns = not significant.**** $p \leq 0.0001$. Throughout figure blue = IB and red = EB.
